# Profiling RNA at chromatin targets in situ by antibody-targeted tagmentation

**DOI:** 10.1101/2022.05.02.490302

**Authors:** Nadiya Khyzha, Steven Henikoff, Kami Ahmad

## Abstract

Whereas techniques to map chromatin-bound proteins are well-developed, mapping chromatin-associated RNAs remains a challenge. Here we describe Reverse Transcribe & Tagment (RT&Tag), in which RNAs associated with a chromatin epitope are targeted by an antibody followed by a protein A-Tn5 transposome. Localized reverse transcription generates RNA/cDNA hybrids that are subsequently tag-mented for sequencing by Tn5. We demonstrate the utility of RT&Tag in Drosophila cells for capturing the noncoding RNA roX2 with the dosage compensation complex and maturing transcripts associated with silencing histone modifications. We also show that RT&Tag can detect N6-methyladenosine (m6A)-modified mRNAs, and show that genes producing methylated transcripts are characterized by extensive promoter pausing of RNA polymerase II. The high efficiency of in situ antibody tethering and tagmentation makes RT&Tag especially suitable for rapid low-cost profiling of chromatin-associated RNAs from small samples.

## Introduction

RNA levels are tightly regulated throughout their lifecycle to ensure proper gene expression^1^. Factors influencing RNA post-transcriptionally include interaction with RNA binding proteins (RBPs), location within the nucleus, and post-transcriptional modifications^1^. The most widely used strategy for assaying these factors is immunoprecipitation, whereby antibodies are used to pull down RNA associated with an epitope of interest from cell lysates^2^. The recovered RNA is then purified and used for downstream applications such as Illumina sequencing^3,4^. Variations of the immunoprecipitation protocol have been developed to study different types of RNA interactions. Examples include RNA immunoprecipitation (RIP) and UV cross-linking and immunoprecipitation (CLIP) for detecting RNA-protein interactions. Chromatin-specific immunoprecipitation assays include Profiling Interacting RNAs on Chromatin followed by deep sequencing (PIRCh-seq) and Chromatin RIP followed by high-throughput sequencing (ChRIP-seq) crosslink RNA to chromatin and assay RNA-chromatin interactions using antibodies targeting histone post-translational modifications^5,6^. Immunoprecipitation assays for N^6^-methyladenosine (m6A) modified RNA include Methylated RNA Immunoprecipitation with next-generation sequencing (MeRIP-seq) and m6A-RIP-seq^7,8^. Unfortunately, these immunoprecipitation-based methods require large sample inputs and optimization of crosslinking condition^2,9^. There is a need for sensitive *in situ* technologies that do not rely on cross-linking or immunoprecipitation to capture endogenous RNA interactions.

Cleavage Under Targets and Tagmentation (CUT&Tag) is an enzyme-tethering strategy developed to profile the binding sites of chromatin proteins within intact nuclei^10^. CUT&Tag bypasses immunoprecipitation and instead uses antibodies to tether a protein A-Tn5 transposase fusion protein *in situ*. Tn5 undergoes a tagmentation reaction where genomic DNA is cleaved and tagged with sequencing adaptors. These sequencing adaptors are then used to generate Illumina sequencing libraries. However, Tn5 also contains an RNase H-like domain that can bind and tagment reverse transcribed RNA/cDNA hybrids^11,12^. This finding inspired us to develop Reverse Transcribe & Tagment (RT&Tag), a proximity labeling tool for capturing RNA interactions within intact nuclei. RT&Tag follows the framework of CUT&Tag but is adapted to capture signal from RNA instead of genomic DNA. Relative to RIP-based immunoprecipitation methods, RT&Tag requires few cells and a low number of sequencing reads, while capturing interactions within intact nuclei. In this work, we demonstrate the general utility of RT&Tag by applying it to a variety of RNA- and chromatin-dependent phenomena in Drosophila S2 nuclei. Specifically, we use RT&Tag to target the dosage compensation complex, Polycomb chromatin domains, and the m6A RNA post-transcriptional modification. Surprisingly, we find that binding of the m6A writer, METTL3, is not sufficient for RNA methylation. Instead, we find that RNA Polymerase II pausing is a strong predictor of m6A mark deposition. This finding illustrates the potential of RT&Tag to empower research in the fields of epigenetics and RNA biology.

## Results

### RT&Tag general workflow

To create a method analogous to CUT&Tag for detecting localized RNAs, we capitalized on the ability of Tn5 to tagment RNA/DNA hybrid duplexes^11,12^. Briefly, we first isolate nuclei, bind a factor-specific primary antibody, bind a streptavidin-conjugated secondary antibody, and then tether biotinylated oligo(dT)-adapter oligos and pA-Tn5 to that (Fig. 1A). Using biotinylated oligo(dT)-adaptor fusions increases the signal-to-noise ratio by selectively priming nearby RNA for reverse transcription (RT) (Supplementary Fig. 1A). Addition of reverse transcriptase then converts mature transcripts near the binding site to RNA/DNA hybrids, which are tagmented by the juxtaposed pA-Tn5. RT and tagmentation are then performed within one incubation step in a compatible buffer. With simultaneous RT and tagmentation, we were able to detect higher transcript enrichment than with sequential RT and tagmentation (Supplementary Fig. 1B). This may be attributed to RT altering RNA secondary structure which could then disrupt RNA-protein interactions or mask epitope binding sites. Hence, the simultaneous RT and tagmentation approach may preserve endogenous RNA interactions until the time of tagmentation without sacrificing RT efficiency (Supplementary Fig. 1C). After RT and tagmentation, the pA-Tn5 is stripped off with SDS and sequencing libraries are amplified using PCR. To generate sequencing libraries only from RNA instead of from genomic DNA, the i7 adaptor sequence is appended 5’ to the oligo(dT) sequence, ensuring its integration into all reverse transcribed transcripts (Supplementary Fig. 2). The i5 adaptor is loaded into Tn5 and is integrated into RNA-cDNA hybrids via tagmentation. As such, only tagmented RNA-cDNA hybrids have both adaptors necessary for library amplification, whereas genomic DNA lacks the i7 adaptor. With the i7 adaptor appended to the oligo(dT), the amplified libraries should detect signal from the 3’ end of the RNA. This means that only a small segment of the RNA needs to be effectively reverse transcribed to be detected by RT&Tag. Not having to reverse transcribe the entirety of the transcripts minimizes variation arising from RT such as interference with the processivity of the reverse transcriptase due to RNA secondary structure, protein binding and RNA length. To explore the capabilities of RT&Tag, we have applied it to address diverse problems in RNA-chromatin biology. These include identifying RNA interacting with proteins and chromatin domains and detection of transcripts enriched for post-transcriptional modifications. We found that all three modalities could be assayed using RT&Tag, unlike immunoprecipitation-based methods which required a different method for each modality (Fig. 1B).

**Fig. 1:**
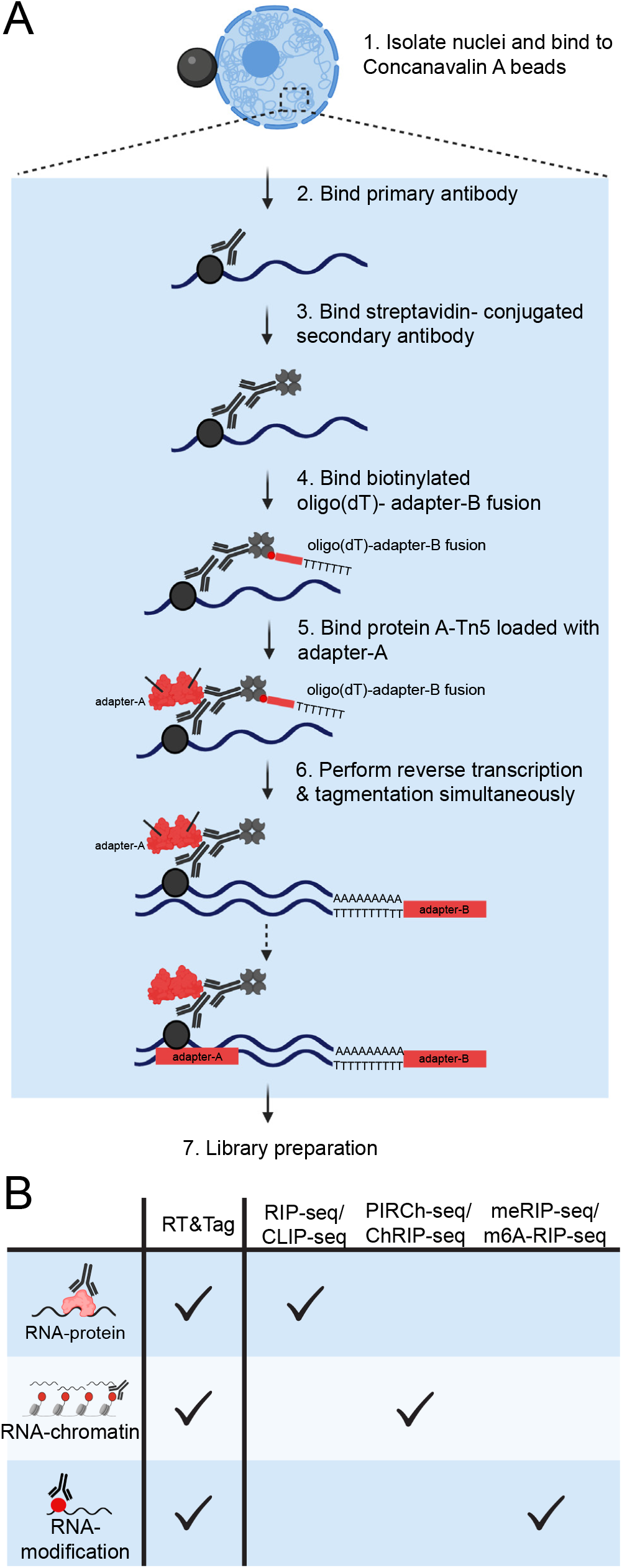
RT&Tag general workflow. **A)** Schematic outlining the steps of RT&Tag: 1) nuclei are isolated and bound to Concanavalin A paramagnetic beads; 2) primary antibody binds epitope of interest; 3) streptavidin-conjugated secondary antibody binds to the primary antibody; 4) biotinylated oligo(dT)-adapter-B fusion oligonucleotide binds to the streptavidin conjugated secondary antibody; 5) protein A-Tn5 loaded with adapter-A binds to the primary and secondary antibodies; 6) simultaneous reverse transcription and tagmentation are then performed to generate RNA-cDNA hybrids that contain the two complementary adaptors; 7) sequencing libraries are amplified using PCR. **B)** Illustration showing applications of RT&Tag described in this work, which include identifying RNA-protein interactions, RNA-chromatin interactions, and RNA post-transcriptional modifications. This is contrasted to immunoprecipitation-based techniques which require a separate method for targeting each type of interaction.

### RT&Tag captures the interaction between MSL2 and the roX2 noncoding RNA

As a proof of concept, we used antibodies to target the RNA-associated dosage compensation complex in the male Drosophila S2 cell line (Fig. 2A). The MSL complex coats the male X chromosome to upregulate gene expression by depositing the activation-associated H4K16ac mark^13^. The long non-coding RNA (lncRNA) roX2 is bound by MSL2, an interaction that we expected to detect using RT&Tag^13^. Using an anti-MSL2 antibody, we generated RT&Tag DNA sequencing libraries. Four features indicated that these libraries resulted from tagmentation of reverse transcribed RNA/DNA hybrids. First, no libraries were produced when reverse transcriptase was omitted (Fig. 2B). Second, while CUT&Tag for chromatin targets produced a nucleosomal ladder, RT&Tag libraries had a broad size distribution ranging predominantly from 200 bp to 1000 bp with no nucleosomal pattern (Fig. 2B). Third, mapped RT&Tag reads were primarily of exonic origin (66%) with a small number of intronic (16%) and intergenic reads (18%) (Fig. 2C, Supplementary Fig. 3A). Finally, reads mostly fell at the 3’ ends of gene bodies consistent with priming from the poly-A tail of mature transcripts by the oligo-dT-adaptor fusion (Fig. 2D, Supplementary Fig. 3B). Altogether, these findings demonstrate that the RT&Tag signal is exclusively from RNA.

**Fig. 2:**
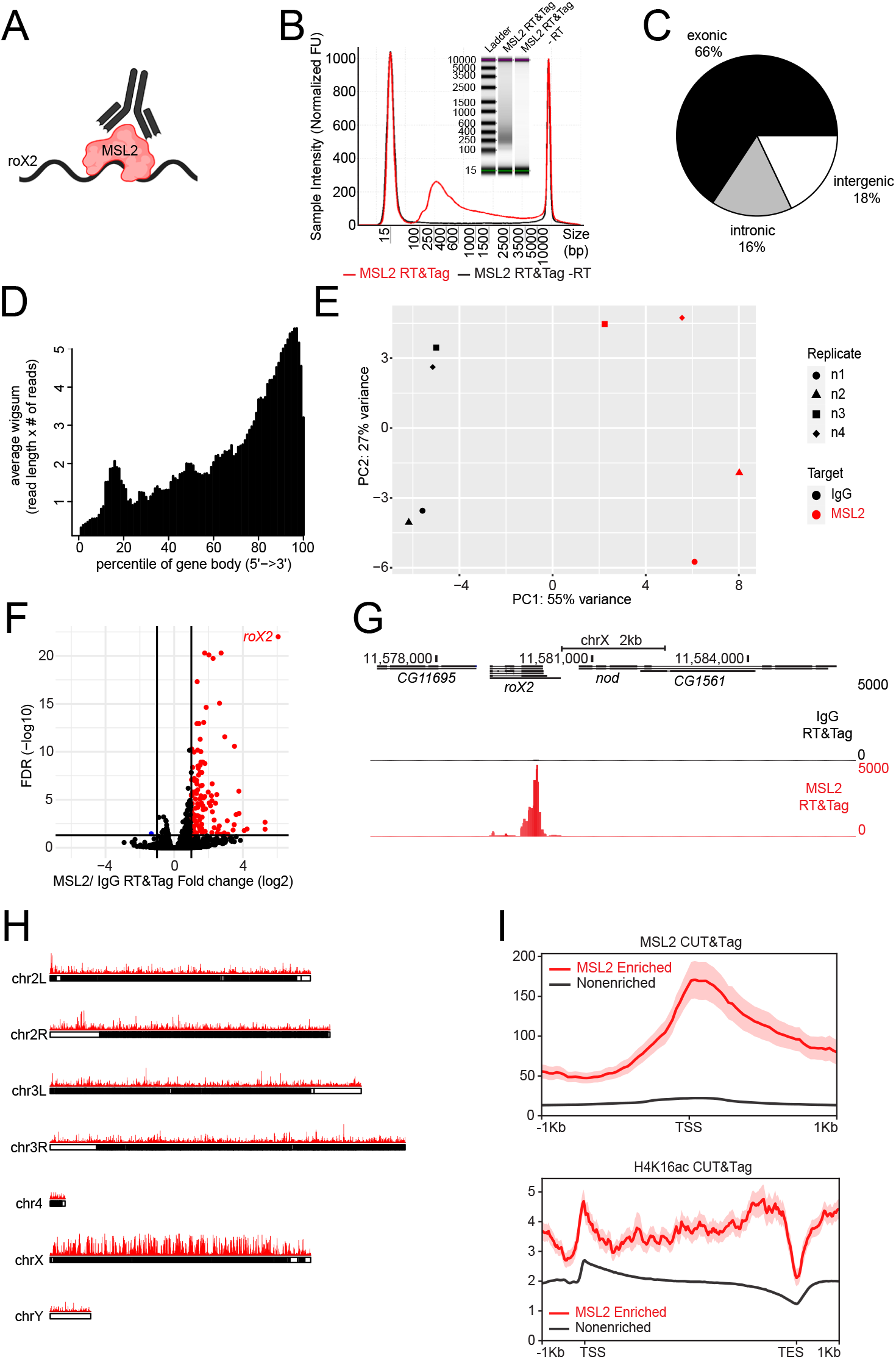
RT&Tag captures the interaction between MSL2 and the *roX2* noncoding RNA. **A)** Illustration showing RT&Tag being used to capture the interaction between MSL2 and *roX2*. **B)** Tapestation gel image and corresponding electropherogram showing size distribution of the MSL2 RT&Tag libraries after two rounds of 0.8x bead clean-up. In the absence of reverse transcriptase (RT), no libraries are produced. **C)** Pie chart showing the proportion of MSL2 RT&Tag reads (n=4) aligning to regions classified as either exonic, intronic, or intergenic. **D)** Density plot showing the distribution of aligned MSL2 RT&Tag reads (n=4) scaled over Drosophila gene bodies. A clear bias towards the 3’ end of genes is observed. A small bump around 15-20^th^ percentile may be explained by internal priming at A-rich stretches, especially those found with the 18S and 26S rRNAs. **E)** Principal component analysis showing separation between IgG and MSL2 RT&Tag samples along the first principal component and separation between replicates in the second principal component. The first two and last two replicates have been sequenced on two separate flow cells and hence a batch effect may be observed. **F)** Volcano plot showing transcripts differentially enriched for MSL2 over IgG RT&Tag (fold change >2, FDR <0.05, n=4). Transcripts enriched for MSL2 are highlighted in red, nonenriched are in black, and depleted are in blue. **G)** Genome browser track showing the distribution of MSL2 and IgG RT&Tag signal over the gene body of *roX2*. Combined reads from 4 replicates are shown. **H)** Karyoplots showing the bins (50 bp) where MSL2 RT&Tag signal is 4-fold over IgG plotted over the Drosophila chromosomes. **I)** Profile plots showing the MSL2 (top) and H4K16ac (bottom) CUT&Tag signal around the TSS (top) and gene bodies (bottom) of MSL2 RT&Tag enriched or nonenriched transcripts.

The performance of MSL2 RT&Tag was then evaluated. Differences between MSL2 RT&Tag and the IgG background control were assessed using principal component analysis (PCA) (Fig. 2E). The first principal component captured a clear separation (55% variance) between IgG and MSL2 libraries. This separation was greater than the second principal component which captured the variability between replicates (27% variance). Differential enrichment of MSL2-targeted transcripts over IgG (>2 Fold Change (FC), <0.05 FDR) identified 121 transcripts, of which roX2 showed very high enrichment and statistical significance (67 FC, <1×10^−22^ FDR; Fig. 2F, Supplementary Table 1). This enrichment of MSL2 RT&Tag signal over IgG is illustrated over the gene body of roX2 using UCSC genome browser tracks, highlighting a clear 3’ bias in the distribution of reads (Fig. 2G). Apart from roX2, 120 transcripts were differentially enriched for MSL2. The MSL2 RT&Tag signal normalized for IgG showed a strong preference for the X-chromosome (56.3% of >4-fold enriched bins, Fig. 2H). Given that MSL2 binds across the *X* chromosome, we asked whether MSL2 RT&Tag captured RNA that was transcribed proximal to these MSL2 binding sites. Hence, we mapped the MSL2 CUT&Tag signal at the transcriptional start sites (TSSs) of MSL2 enriched or nonenriched transcripts. Additionally, H4K16ac CUT&Tag signal was mapped over the gene bodies of MSL2 enriched or nonenriched transcripts. Higher MSL2 and H4K16ac CUT&Tag signal was observed for MSL2 RT&Tag enriched than non-enriched transcripts, supporting our hypothesis (Fig. 2I). Furthermore, 75% of MSL2-enriched transcripts were within 13 kb of an MSL2 binding peak which is much closer than for nonenriched transcripts (12,608 bp vs 2,841,851 bp, p< 2.2×10^−16^, Supplementary Fig. 4A). As an example, MSL2 and H4K16ac CUT&Tag signal can be seen over the gene bodies of MSL2 RT&Tag enriched transcripts, *ph-d* and *pcx* (Supplementary Fig. 4B). Overall, these results show that RT&Tag recapitulates the well-known MSL2-roX2 interaction and captures interactions between MSL2 and transcripts found within its vicinity. Distinguishing direct versus proximal interactions may be guided by the idea that proximity interactions should be transient and result in weaker enrichment. As such, enrichment for roX2 is a unique outlier both in fold change and FDR while the proximal transcripts found on the X-chromosome exhibit either low fold change or low FDR.

We then compared our MSL2 RT&Tag data to a published RIP-seq dataset, which targeted a subunit of the Drosophila MSL complex, maleless (MLE). Like RT&Tag, MLE RIP-seq was able to identify the interaction between MLE and roX2 in S2 cells (Supplementary Fig. 5A). However, to achieve a comparable degree of enrichment for roX2, RIP-seq required 500 times the number of cells and 4 times as many sequencing reads as RT&Tag (Supplementary Fig. 5B). Apart from the roX RNAs, RT&Tag and RIP-seq picked up transcripts that were unique to each method (Supplementary Fig. 5C). Transcripts unique to RT&Tag were predominantly transcribed from the X-chromosome unlike the transcripts unique to RIP-seq (Supplementary Fig. 5D). This comparison highlights the fundamental difference between RT&Tag and immunoprecipitation-based methods. Being a proximity labelling technique, RT&Tag can pick up transcripts near MSL complex binding sites, whereas RIP-seq captures binding interactions within cell lysates, some of which might not occur under endogenous conditions.

### RT&Tag captures transcripts within Polycomb domains

After validating RT&Tag using MSL2, we applied RT&Tag to identify RNA associated with chromatin domains (Fig. 3A). Polycomb domains are large regions of chromatin decorated with repressive histone H3K27me3 marks^14,15^. They make for an appealing target as studies in mammals have implicated RNA in their establishment and maintenance^15^. Targeting H3K27me3 with an antibody, RT&Tag identified 1,342 transcripts that are differentially enriched for H3K27me3 over IgG background (>2 FC, <0.05 FDR; Fig. 3B, Supplementary Table 2). As examples, the H3K27me3-targeted RT&Tag signals are shown over the two most statistically significant hits, the lncRNAs *CR43334* and *CR42862* (Fig. 3C). We then assessed the performance of H3K27me3 RT&Tag with decreasing numbers of input nuclei. The H3K27me3 RT&Tag signal was highly reproducible using 100,000 and 25,000 nuclei (Supplementary Fig. 6A) and even 5,000 nuclei for *CR43334* and *CR42862* (Supplementary Fig. 6B). We then proceeded to characterize H3K27me3-enriched transcripts and found them to be predominantly protein coding (1,178 out of 1,342) with low expression levels (mean 16.6 counts per million (CPM) vs 97.1 CPM for nonenriched genes, p<2.2×10^−16^) (Fig. 3D, E). Additionally, H3K27me3 RT&Tag-enriched transcripts had more repressive H3K27me3 CUT&Tag signal and lower active H3K36me3 and H3K4me3 CUT&Tag signal at their TSS or over their gene bodies than nonenriched transcripts (Fig. 3F). In line with this, H3K27me3 RT&Tag-enriched transcripts enriched for GO terms associated with developmental biological processes, which are associated with Polycomb^16^ (Supplementary Fig. 7A). Altogether, these data suggest that H3K27me3 RT&Tag-enriched transcripts are from repressed genes within Polycomb domains. These include classic examples of Polycomb repressed genes such as the Hox genes^17^, which we have found to show strong enrichment for H3K27me3-targeted RT&Tag signal (Fig. 3G, Supplementary Fig. 6C).

**Fig. 3:**
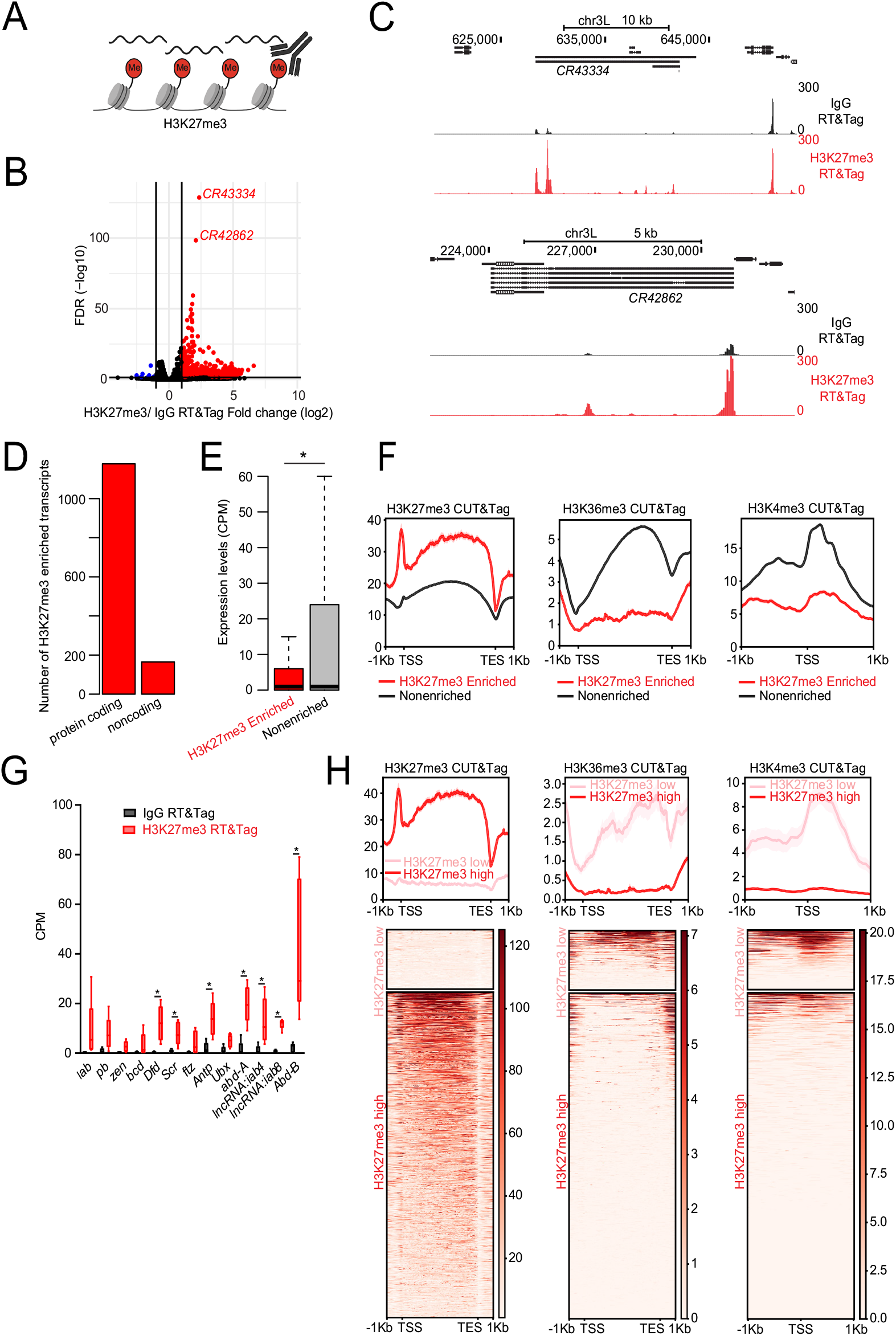
RT&Tag captures transcripts within Polycomb domains. **A)** Illustration showing RT&Tag being used to capture transcripts within H3K27me3-demarcated Polycomb domains. **B)** Volcano plot showing transcripts that are differentially enriched for H3K27me3 RT&Tag over IgG (fold change >2, FDR <0.05, n=5). Genes enriched for H3K27me3 are highlighted in red, nonenriched are in black, and depleted are in blue. The two most highly significant transcripts are labelled. **C)** Genome browser track showing the distribution of H3K27me3 and IgG RT&Tag signal over the gene bodies of *CR43334* and *CR42862*. Combined reads from 5 replicates are shown. **D)** Bar graph showing the number of H3K27me3 enriched transcripts that are protein coding or noncoding. **E)** Boxplot showing the RNA-seq expression levels (Counts per million, CPM) of H3K27me3 enriched or nonenriched transcripts. *p<0.05, unpaired t-test, n=5. **F)** Profile plots showing the H3K27me3 (left), H3K36me3 (middle) and H3K4me3 (right) CUT&Tag signal around the gene bodies or TSS of genes that were categorized as being enriched for H3K27me3 RT&Tag or nonenriched. **G)** Boxplots showing the IgG and H3K27me3 RT&Tag signal (Counts per million, CPM) for the HOX cluster genes. *FDR<0.05, n=5. **H)** Profile plots and heatmaps showing the H3K27me3 (left), H3K36me3 (center) and H3K4me3 (right) signal over the gene bodies or TSS of H3K27me3 RT&Tag enriched transcripts that have high or low levels of H3K27me3 CUT&Tag signal over their gene bodies. Heatmaps are plotted in order of decreasing H3K27me3 (left), H3K36me3 (center) and H3K4me3 (right) CUT&Tag signal.

We then assessed what proportion of H3K27me3-targeted RT&Tag transcripts were transcribed from regions decorated by H3K27me3 marks. First, we established the H3K27me3 CUT&Tag background level cut-off in S2 cells as the H3K27me3 CUT&Tag signal over the gene bodies for the top 25% expressed genes (>17 CPM) (Supplementary Fig. 7B). Using this cut-off, 84.5% (1,134 out of 1,342) of H3K27me3-RT&Tag enriched transcripts were found to be from regions with substantial H3K27me3 CUT&Tag signal (Fig. 3H). These genes also show low levels of active H3K36me3 and H3K4me3 CUT&Tag signal (Fig. 3H). The remaining 208 H3K27me3-directed RT&Tag enriched tran-scripts are from outside of H3K27me3 marked regions and show high H3K36me3 and H3K4me3 CUT&Tag signals. These 208 H3K27me3 RT&Tag-enriched genes are more highly expressed than those from H3K27me3 marked regions (mean 60.9 vs 8.5 CPM, p<0.004; Supplementary Fig. 7C). Given that transcripts captured by RT&Tag must have poly(A) tails, our findings are consistent with the low production of new transcripts from silenced regions, and the subsequent capture of these transcripts near their sites of transcription^18,19^.

### RT&Tag captures transcripts enriched for the m6A post-transcriptional modification

Having demonstrated that RT&Tag can detect RNAs in protein complexes, and chromatin domains, we tested if our method could be used for RNA modifications. N^6^-Methyladenosine (m6A) is the most abundant mRNA post-transcriptional modification and has been implicated in numerous aspects of RNA metabolism^20^. Commercial antibodies targeting m6A are available and have been used in RNA immunoprecipitation-based methods (i.e., MeRIP-seq and m6A-seq)^7,8^. Although these techniques are valuable for pinpointing the location of m6A modifications, they require large amounts of input material and suffer from low reproducibility^21^. We reasoned that RT&Tag could provide insights into whether a particular transcript is enriched or depleted for m6A relative to IgG input control (Fig. 4A). Using RT&Tag, we identified 281 transcripts enriched for m6A (>1.5 FC, <0.05 FDR) and 106 transcripts depleted for this modification (>1.5 FC, <0.05 FDR; Fig. 4B, Supplementary Table 3). Of these, *aqz, Syx1A, gish, pum* and *Prosap* transcripts have been previously reported as enriched for m6A (Ref. 22, Fig. 4B, C). Next, we assessed the performance of m6A RT&Tag with varying numbers of input nuclei. The m6A RT&Tag signal was highly reproducible using 100,000 and 25,000 nuclei (Supplementary Fig. 8A) and even 5,000 nuclei for *aqz* and *Syx1A* (Supplementary Fig. 8B). Transcripts enriched for m6A are associated with development and transcription factor binding Gene Ontology (GO) terms, whereas transcripts depleted for m6A tend to be associated with housekeeping GO terms, especially translational components and processes (Fig. 4D).

**Fig. 4:**
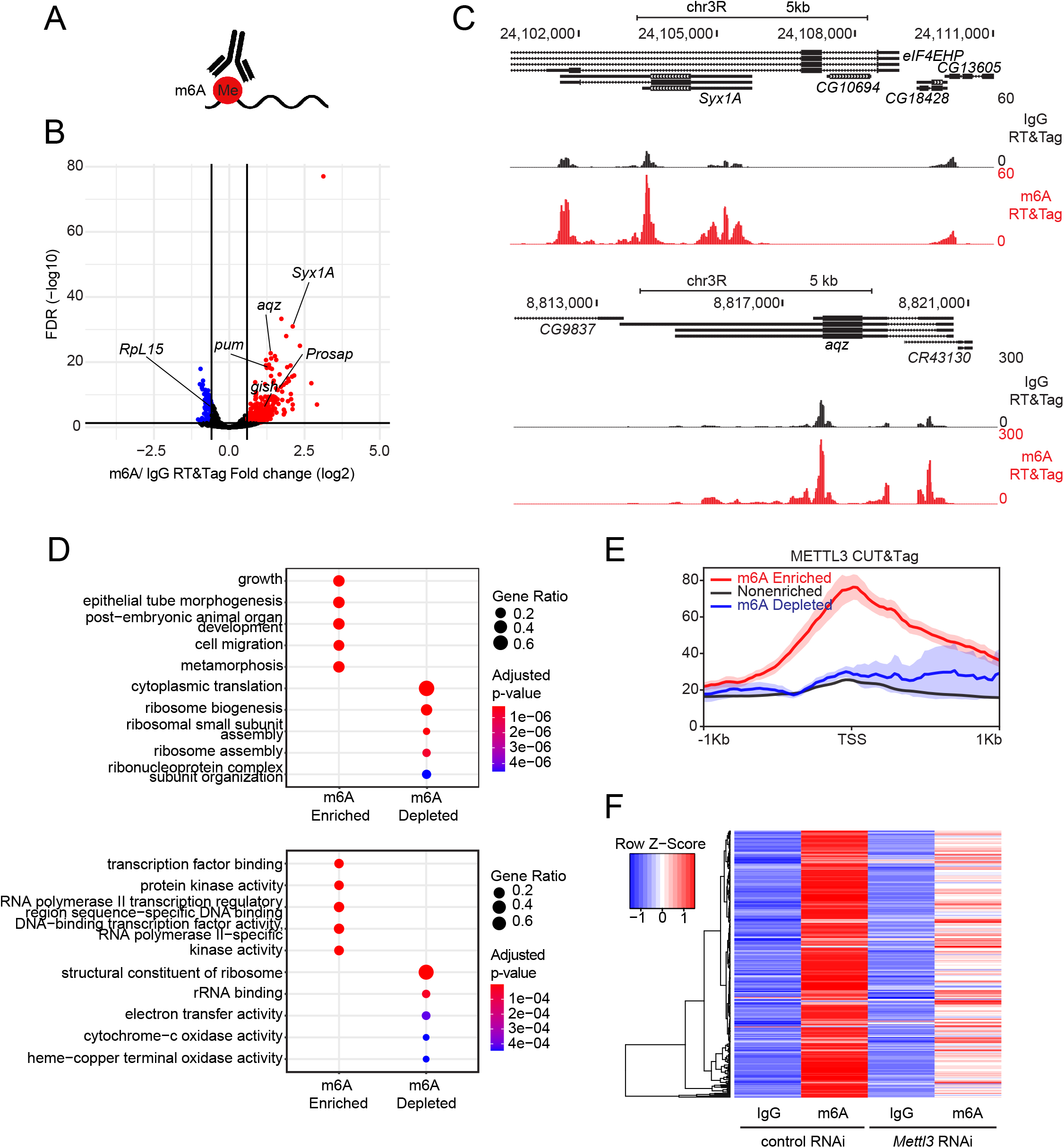
RT&Tag captures transcripts enriched for the m6A posttranscriptional modification. **A)** Illustration showing RT&Tag being used to capture transcripts enriched for the m6A post-transcriptional modification. **B)** Volcano plot showing genes that are differentially enriched for m6A over IgG RT&Tag (fold change >1.5, FDR <0.05, n=3). Genes enriched for m6A are highlighted in red, nonenriched are in black, and m6A depleted are in blue. Genes previously shown to be enriched or depleted for m6A are labelled. **C)** Genome browser track showing the distribution of m6A and IgG RT&Tag reads over the gene bodies of *aqz* and *Syx1A*. Combined reads from 3 replicates are shown. **D)** Dot plot showing the top 5 GO biological process (top) and molecular function (bottom) terms associated with m6A enriched and m6A depleted transcripts. The dot size corresponds to the gene ratio (# genes related to GO term / total number of m6A enriched or depleted genes) and the color represents statistical significance. **E)** Profile plots showing the METTL3 CUT&Tag signal at the TSS of genes that are enriched, nonenriched, or depleted for m6A. **F)** Heatmap showing IgG or m6A RT&Tag counts for m6A enriched genes in S2 cells treated with either control or *Mettl3* RNAi. Values are scaled based on rows.

The Drosophila homologue of the METTL3 methyltransferase binds to chromatin and catalyzes the m6A modification on nascent transcripts^23^. We observed high levels of METTL3 CUT&Tag signal at the TSSs of m6A enriched genes, relative to nonenriched or m6A depleted genes (Fig. 4E). To validate our list of m6A enriched genes, we knocked down the gene encoding METTL3 (*Mettl3*, formerly called *Inducer of meiosis* in yeast or *Ime4*) levels by 80% using RNAi (Supplementary Fig. 9A). Doing so resulted in a modest decrease (>10%) for 81% of m6A enriched transcripts (Fig. 4F). Altogether, these results show that m6A enriched transcripts identified by RT&Tag are METTL3 methylation dependent.

### Genes of methylated transcripts are characterized by promoter proximally paused RNA Polymerase II

Whereas the promoters of genes producing m6A-enriched transcripts are enriched for METTL3, we noticed that the METTL3 CUT&Tag signal at TSSs of m6A-depleted transcripts was still above IgG CUT&Tag signal (Supplementary Fig. 9B). In fact, METTL3 binding was widely observed amongst the top 25% expressed genes (>17CPM) (Fig. 5A). Indeed, total RNA polymerase II (RNAPolII) and METTL3 binding are positively correlated (Fig. 5A, Supplementary Fig. 9C)^24,25^. Thus, we reasoned that METTL3 must be preferentially recruited to sites of active transcription. This leads to the expectation that highly expressed transcripts would be enriched for transcript methylation. However, m6A-enriched transcripts tend to be expressed at lower levels than m6A-depleted transcripts (192 CPM vs 4,856 CPM, p=6.2×10^−7^; Fig. 5B). In line with expression level differences, genes producing m6A-enriched transcripts have lower levels of active H3K4me3 and H3K36me3 marks (Fig. 5C). Hence, the m6A methylation mark is not associated with high level of transcription. We then asked whether increasing METTL3 levels at a gene would in turn result in more transcript methylation. Heat shock of Drosophila cells induces a large influx of RNAPolII into the bodies of heat shock protein (HSP) genes^26^, which we can observe by CUT&Tag (Fig. 5D). In addition to RNAPolII enrichment, we found that heat shock causes a dramatic increase in METTL3 (Fig. 5D). This increase is not limited to promoters, but now extends into the bodies of the *Hsp70* genes. However, induced *Hsp70* transcripts do not accumulate the m6A modification, despite the large influx of METTL3 and presence of RRACH motifs (the RNA sequence in which the m6A modification occurs) within the *Hsp70* transcripts (Fig. 5E, Supplementary Fig. 9D). Thus, METTL3 binding on its own does not reliably predict methylation status.

**Fig. 5:**
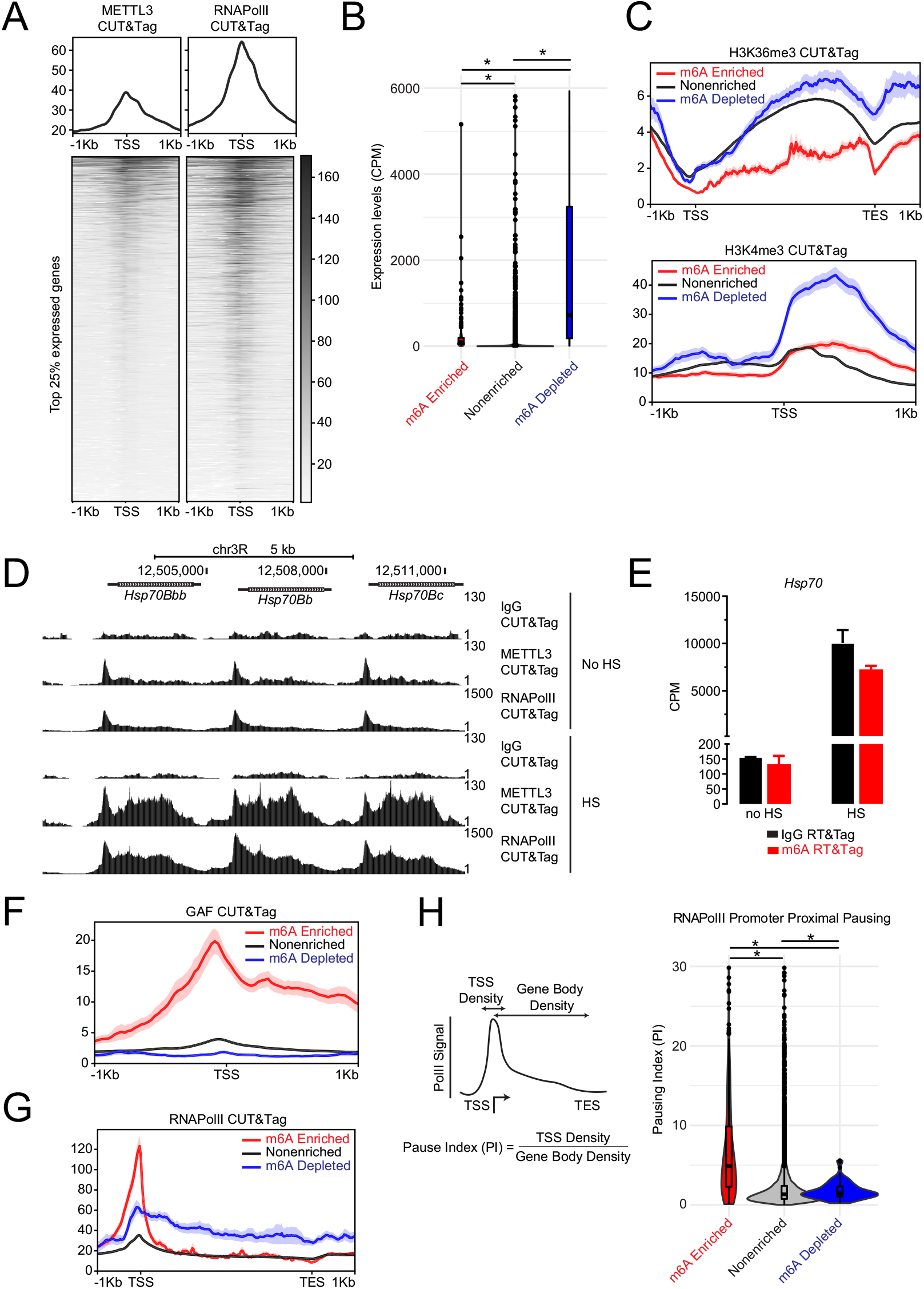
Genes of methylated transcripts are characterized by promoter proximally paused RNA Polymerase II. **A)** Profile plots and heatmaps showing METTL3 (left) and total RNA polymerase II (RNAPolII, right) CUT&Tag signal at the TSS of the top 25% expressed genes. Heatmaps are plotted in the order of decreasing METTL3 signal. **B)** Violin plots showing the RNA-seq expression levels (counts per million, CPM) of genes that are depleted, enriched, or nonenriched for m6A. *p<0.05, unpaired t-test. **C)** Profile plots showing the H3K36me3 (top) and H3K4me3 (bottom) CUT&Tag signal over the gene bodies or at the TSS of genes that are enriched, non-enriched, or depleted for m6A. **D)** Genome browser tracks showing the IgG, METTL3, and RNAPolII CUT&Tag signal over the gene bodies of *Hsp70* genes with no heat shock (no HS) or after 15 minutes of heat shock (HS). **E)** Bar graph showing the IgG and m6A RT&Tag signal for *Hsp70* with no heat shock (no HS) and after 15 minutes of heat shock (HS). **F)** Profile plot showing GAGA factor (GAF) CUT&Tag signal at the TSS of m6A enriched, nonenriched, and depleted transcripts. **G)** Profile plot showing RNAPolII CUT&Tag signal over the gene bodies of m6A enriched, nonenriched, and depleted transcripts. **H)** Schematic showing how the promoter proximal pausing index (PI) was calculated (left). PI was calculated by dividing the promoter (+/- 250 bp around the TSS) RNAPolII CUT&Tag signal over the gene body RNAPolII CUT&Tag signal. Violin plots displaying the PI of m6A enriched, nonenriched, and depleted transcripts (right). *p<0.05, unpaired t-test.

What other features might distinguish m6A-enriched and depleted transcripts? Motif analysis revealed GAGA motifs within the promoters of m6A-enriched transcripts (Supplementary Fig. 9E). GAGA factor (GAF) is a DNA-binding transcription factor that binds GAGA motifs and is associated with promoter proximal pausing of RNAPolII^27^. In line with GAGA motif enrichment, much higher GAF CUT&Tag signal is detected at the TSSs of m6A-enriched (Fig. 5F). For this reason, we looked at the distribution of total RNAPolII signal over gene bodies relative to the TSS. We observed m6A-enriched transcripts to have more RNAPolII signal at the TSS and less within gene bodies (Fig. 5G). We then calculated the RNAPolII promoter proximal Pausing Index (PI) as the ratio of RNAPolII signal at the promoter (±250 bp around the TSS) to signal over the gene body. Indeed, m6A-enriched transcripts had very high levels of PI relative to m6A-depleted transcripts (7.4 vs 1.8, p< 2.2e^-16^) (Fig. 5H). This high level of PI was not related to the expression level of the m6A-enriched transcripts (Supplementary Fig. 9F). Altogether, our findings suggest that transcripts with a very high degree of polymerase pausing and high GAF binding at their promoters are predominantly enriched for the m6A post-transcriptional modification.

## Discussion

In this work we developed RT&Tag, a proximity labeling tool, that uses antibodies to tether Tn5 and tagment near-by RNA within intact nuclei. RT&Tag fundamentally differs from immunoprecipitation-based methods which capture RNA binding to factors within a cell lysate instead of endogenous proximity interactions. Furthermore, RT&Tag does not require cross-linking or RNA fragmentation, and the same RT&Tag protocol can be applied to RNA-protein interactions, RNA-chromatin interactions, and RNA modifications. In contrast, immunoprecipitation techniques require separate protocols for each application.

A major advantage of RT&Tag over immunoprecipitation is its efficiency. RT&Tag requires fewer than ∼100,000 cells which is at least 50-fold fewer than the number needed for PIRCh-seq and ChRIP-seq (Table 1)^5,6^. RT&Tag also works with few sequencing reads as the RT&Tag reads are concentrated at the 3’ end of RNA^28^. Specifically, we have had success with 4-8 million reads per sample for RT&Tag, relative to PIRCh-seq where around 50 million reads were used (Table 1)^5^. Other enzyme tethering based techniques are emerging as in situ alternatives to immunoprecipitation. For example, APEX sequencing (APEX-seq) and Targets of RNA-binding proteins Identified By Editing (TRIBE) tether RNA modifying enzymes by fusing them with other proteins^29-31^. However, these methods have yet to be used to identify RNA interactions occurring on chromatin. Additionally, the need to generate fusion proteins for each protein target makes these techniques laborious and low throughput, unlike RT&Tag, which can be easily applied to any epitope with an available antibody. Another advantage of RT&Tag is that RNA/cDNA hybrids are directly tagmented by Tn5 with sequencing adaptors. This allows for seamless generation of Illumina sequencing libraries using a simple PCR reaction, without the need to purify RNA as in ChRIP-seq, APEX-seq and TRIBE. The lack of purification steps makes RT&Tag adaptable for automation as was done with AutoCUT&Tag^32^. Together with low cell number input, low sequencing depth, RT&Tag presents a high throughput method to study RNA metabolism by targeting chromatin factors and post-translational modifications.

**Table 1:**
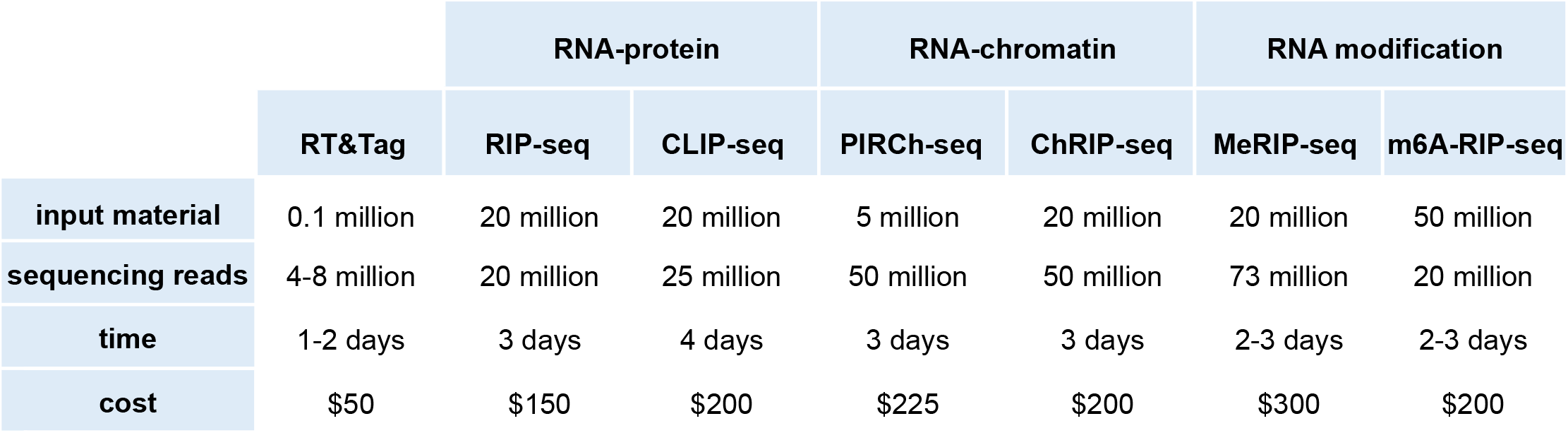
Comparison of RT&Tag to immunoprecipitation-based methods.

Using RT&Tag, we gained insight into the N-methyladenosine (m6A) modification. m6A is the most prevalent mRNA post-transcriptional modification and has been implicated in splicing, mRNA decay, and translation^20^. The m6A modification is catalyzed by the methyltransferase, METTL3^23^. How METTL3 discriminates which RNAs get methylated is unclear. We have observed widespread METTL3 binding at the promoters of expressed genes. However, we found that most of these genes were not enriched for m6A, suggesting that other factors must be involved. Instead, we found RNAPolII promoter pausing to be a strong predictor of m6A deposition. We were surprised that *Hsp70*, a gene known to exhibit RNAPolII pausing, was not identified as being m6A-enriched using RT&Tag. However, upon calculating the pausing index of *Hsp70*, we have found it to be on par with that of m6A nonenriched transcripts. This suggests that only genes exhibiting very high levels of RNAPolII pausing are enriched for m6A. RNAPolII dynamics, especially elongation speed, have previously been implicated in regulating co-transcriptional processes including splicing and alternative polyadenylation^33^. Furthermore, human MCF7 breast cancer cells expressing a slow elongation RNAPolII mutant have been reported to have increased m6A levels^34^. How RNAPolII promoter pausing contributes to m6A deposition is not known but may be due to the increased amount of time METTL3 is bound near the promoter. As such, METTL3 would have more contact time with the 5’ end of RNA, the region where m6A is predominantly found in Drosophila^35^. METTL3 itself has been found to promote productive RNAPolII elongation, which suggests that there may be two-way communication between m6A and RNAPolII processivity^25,36^. An alternative explanation for the discrepancy between METTL3 binding and m6A levels is that methylation may occur at all METTL3 bound transcripts but not be retained. Fat mass and obesity-associated protein (FTO) is a demethylase that is known to remove the m6A mark after transcription in mammals. However, no FTO homologue has been identified in Drosophila^23^. Deposition of m6A at splice junctions and introns of nascent transcripts has been implicated in regulating splicing^37^. Thus, intronic m6A marks may be lost during splicing and not be captured by m6A RT&Tag, which specifically measures m6A levels in mature transcripts. Altogether, our findings suggest METTL3 binding does not correspond to the presence of m6A, and that additional factors are necessary for transcript methylation.

Being a proximity tagmentation tool, RT&Tag could have numerous applications given there is an available antibody. Although this work described only chromatin applications, RT&Tag is not necessarily limited to chromatin, and future studies might adapt RT&Tag for targets in the cytoplasm, such as RNA-protein interactions and RNA post-transcriptional modifications. Efforts to catalogue RNA binding protein (RBP) bound transcripts are still in their infancy. Phase 3 of the ENCODE consortium profiled 150 RBPs using immunoprecipitation in the HepG2 and K562 cell lines^38^. Given that the human genome contains over 1500 RBP-encoding genes and mutations in RBPs are becoming implicated in genetic diseases, much work remains to be done to characterize their bound transcripts^39,40^. Similarly, cataloguing sites of m6A modification on a large scale is yet to be done. METTL3 knock-out experiments in mammals (human and mice) have shown that m6A is required for cell differentiation and embryonic viability^41-44^. The commonly used MeRIP-seq and m6A-seq techniques require large amounts of RNA input which makes them impractical for studying differentiating cells and development. RT&Tag can fill the need for high throughput profiling of chromatin-bound RBP-RNA interactions and m6A enriched transcripts, especially when sample input is limiting such as with clinical samples or embryonic cells.

## Methods

### Cell culture and nuclei preparation

Drosophila S2 cells were obtained from Invitrogen (10831-014) and were cultured in HyClone SFX-Insect cell culture media (HyClone) supplemented with 18 mM L-Glutamine (Sigma-Aldrich). S2 cells were maintained at the confluency of 2-10 million cells per mL at 25°C. To induce the heat shock response, S2 cells were placed at 37°C for 15 minutes. To prepare nuclei for CUT&Tag and RT&Tag, 4 million S2 cells were collected by centrifuging at 300 g for 5 minutes followed by a wash with 1x PBS. Nuclei were then isolated by incubating with NE1 buffer (10 mM HEPES pH7.9, 10 mM KCl, 0.1% Triton X-100, 20% glycerol, 0.5 mM spermidine, Roche Complete Protease Inhibitor Cocktail) for 10 minutes on ice. The nuclei were then centrifuged at 500 g for 8 minutes and resuspended in Wash Buffer (20 mM HEPES pH7.5, 150 mM NaCl, 0.5 mM spermidine, Roche Complete Protease Inhibitor Cocktail). The nuclei were either used fresh or were frozen in Wash Buffer with 10% DMSO and stored at -80°C. For RT&Tag, the NE1 and Wash buffers were supplemented with 1 U/μL of RNasin Ribonuclease Inhibitor (Promega).

### Antibodies

The following primary antibodies were used for RT&Tag and CUT&Tag experiments: rabbit anti-IgG (Abcam ab172730), rabbit anti-MSL2 (gift from Mitzi Kuroda, Harvard Medical School), rabbit anti-H4K16ac (Abcam ab109463), rabbit anti-H3K27me3 (Cell Signaling Technology CST9733), rabbit anti-H3K36me3 (Thermo MA5-24687), rabbit anti-H3K4me3 (Thermo 711958), rabbit anti-m6A (Megabase AP60500), rabbit anti-METTL3 (Proteintech 15073-1-AP), mouse anti-unphosphorylated RNA polymerase II (Abcam ab817), rabbit anti-GAF (gift from Giovanni Cavalli, CNRS Montpellier France). The following secondary antibodies were used: Guinea Pig anti-Rabbit (Antibodies Online ABIN101961) and Rabbit anti-Mouse (Abcam ab46450). Streptavidin conjugated secondary antibodies were generated using the Streptavidin Conjugation Kit (Abcam ab102921) as per manufacturer’s instructions.

### RT&Tag

Single loaded pA-Tn5 was assembled prior to starting RT&Tag. First, the Mosaic end-adapter A (ME-A) and its reverse (ME-Rev) oligonucleotides were annealed in Annealing Buffer (10 mM Tris pH8, 50 mM NaCl, 1 mM EDTA) by heating them at 95 °C for 5 minutes and slowly allowing them to cool to room temperature (Supplementary Table 4). Afterwards, 16 µL of 100 µM annealed ME-A were mixed with 100 µL of 5.5 µM pA-Tn5 for 1 hour at room temperature and stored at -20 °C for future use. S2 nuclei were isolated and bound to paramagnetic Concanavalin A (ConA) beads (Bangs Laboratories). To do so, ConA beads were first activated via 2 washes with Binding Buffer (10 mM HEPES pH7.9, 10 mM KCl, 1 mM CaCl_2_, 1 mM MnCl_2_). Afterwards, 100,000 S2 nuclei were bound to 5 μL of ConA beads for 10 minutes at room temperature. The ConA bound nuclei were then incubated with primary antibody diluted 1:100 in Antibody Buffer (20 mM HEPES pH7.5, 150 mM NaCl, 0.5 mM spermidine, Roche Complete Protease Inhibitor Cocktail, 2mM EDTA, 0.1% BSA and 1 U/μL RNasin Ribonuclease Inhibitor) at 4°C overnight. Afterwards, nuclei were incubated with streptavidin conjugated secondary antibody diluted 1:100 in Wash Buffer (20 mM HEPES pH7.5, 150 mM NaCl, 0.5 mM spermidine, Roche Complete Protease Inhibitor Cocktail) for 45 minutes at RT. Two rounds of washes with Wash Buffer were then performed and nuclei were incubated with 0.2 mM biotinylated oligo(dT)-ME-B in Wash Buffer for 20 minutes at RT. Two rounds of washes with Wash Buffer were then performed and nuclei were incubated with ME-A loaded pA-Tn5 diluted 1:200 in 300 Wash Buffer (20 mM HEPES pH7.5, 300 mM NaCl, 0.5 mM spermidine, Roche Complete Protease Inhibitor Cocktail, and 1 U/μL RNasin Ribonuclease Inhibitor) for 1 hour at RT. ConA bound nuclei were then washed thrice with 300 Wash Buffer. Simultaneous reverse transcription and tagmentation were then performed by resuspending nuclei in MgCl_2_ containing Reverse Transcription buffer (1x Maxima RT Buffer which contains 50 mM Tris-HCl pH 8.3, 75 mM KCl, 3 mM MgCl_2_, 10 mM DTT along with, 0.5 mM dNTPs, 10 U/μL of Maxima H Minus Reverse Transcriptase, and 1 U/μL of RNasin Ribonuclease Inhibitor) for 2 hours at 37°C. The nuclei were then washed with 10 mM TAPS and pA-Tn5 was stripped off by resuspending nuclei in 5 μL of Stripping Buffer (10 mM TAPS with 0.1% SDS) and incubating for 1 hour at 58°C. Libraries were then generated using PCR. The nuclei suspension was mixed with 15 μL of 0.67% Triton X-100, 2 μL of 10 mM i7 primer, 2 μL of 10 mM i5 primer and 25 μL of 2x NEBNext Master Mix (NEB). The following PCR conditions were used: 1) 58°C for 5 minutes, 2) 72°C for 5 minutes, 3) 98°C for 30 seconds, 4) 98°C for 10 seconds, 5) 60°C for 15 seconds, 6) Repeat steps 4-5 13 times, 7) 72°C for 2 minutes, 8) Hold at 4°C. Sequencing libraries were then purified using 0.8x HighPrep PCR Cleanup System (MagBio) beads as per manufacturer’s instructions. Libraries were then resuspended in 21 μL of 10 mM Tris-HCl pH8. Library concentrations were quantified using the High Sensitivity D5000 TapeStation system (Agilent).

### CUT&Tag

CUT&Tag was carried out as described previously (dx.doi.org/10.17504/protocols.io.bqwvmxe6)^10^. Briefly, S2 nuclei were bound to ConA beads at the ratio of 100,000 nuclei per 5 μL beads for 10 minutes at room temperature.

Nuclei were then incubated with primary antibody (1:100) at 4°C overnight followed by secondary antibody (1:100) for 45 minutes at RT the next day. Excess antibody was removed via 2 rounds of washes and the nuclei were incubated with loaded pA-Tn5 (1:200) for 1 hour at RT. Nuclei were washed thrice to remove excess pA-Tn5 and then MgCl_2_ was added to perform tagmentation for 1 hour at 37°C. The reaction was then stopped by doing a wash with 10 mM TAPS and stripping off pA-Tn5 by resuspending nuclei in 0.1% SDS buffer and incubating for 1 hour at 58°C. The SDS was then neutralized with Triton X-100 and libraries were amplified with NEBNext Master Mix (NEB) using 12 rounds of amplification. Sequencing libraries were then purified using 1.2x ratio of HighPrep PCR Cleanup System (MagBio) as per manufacturer’s instructions. Libraries were then resuspended in 21 μL of 10 mM Tris-HCl pH8. Library concentrations were quantified using the D1000 TapeStation system (Agilent).

### RNA interference (RNAi)

PCR templates for in vitro transcription (IVT) were amplified from S2 cell cDNA or pGF-P5(S65T) plasmid using Phusion Hot Start Flex DNA Polymerase (NEB) and primers listed in Supplementary Table 5. PCR products were purified using NucleoSpin® Gel and PCR Clean-Up Kit (Clontech). IVT was performed to generate dsRNA using the T7 High Yield RNA Synthesis Kit (NEB). Template DNA was removed using Turbo DNAse (Ambion) and dsRNA was purified using the NucleoSpin® RNA Cleanup XS kit (Clontech). To perform RNAi, S2 cells were seeded at a density of 1 million cells/ mL of serum-free medium. As control RNAi, a total of 30 μg of GFP dsRNA was added to cells. For *Mettl3* RNAi, 15 μg of *Mettl3* dsRNA #1 plus 15 μg of *Mettl3* dsRNA #2 were added. After 6 hours, medium was replaced with serum containing medium. Treatment with dsRNA was repeated after 48 and 96 hours. Cells were collected after 120 hours.

### RT-qPCR

Total RNA was extracted from S2 cells using the RNeasy Plus Mini Kit (Qiagen) according to manufacturer’s instructions. cDNA was synthesized using the Maxima H Minus Reverse Transcriptase (Thermo Scientific). Real time PCR was performed with the Maxima™ SYBR™ Green qPCR Master Mix (Thermo Scientific) using the ABI QuantStudio5 Real Time PCR Systems instrument. Primers used are listed in Supplementary Table 6. Gene expression levels were quantified using the delta delta C^t^ method using the *Ribosomal Protein L32* (*RPL32*) gene for normalization.

### RNA-sequencing

Total RNA from S2 cells was isolated using the RNeasy Plus Mini Kit (Qiagen). Maxima H Minus Reverse Transcriptase (Thermo Fisher Scientific) was used as per manufacturer’s instructions for first strand synthesis. Reverse transcription was primed using the oligo(dT)-ME-B fusion oligonucleotide. Tagmentation was then performed using 100 ng of RNA-cDNA hybrids, ME-A loaded pA-Tn5, and tagmentation buffer (20 mM HEPES pH7.5, 150 mM NaCl, 10 mM MgCl_2_) for 1 hour at 37°C. Tagmented RNA-cDNA hybrids were purified using 1x ratio of HighPrep PCR Cleanup System (MagBio) as per manufacturer’s instructions. Sequencing libraries were then amplified using NEBNext Master Mix (NEB) using 12 cycles. Libraries were then purified using 0.8x ratio of HighPrep PCR Cleanup System (MagBio) as per manufacturer’s instructions. Libraries were then resuspended in 21 μL of 10 mM Tris-HCl pH8 and quantified using the D5000 TapeStation system (Agilent).

### Sequencing and data preprocessing

For RT&Tag and RNA-sequencing, single-end 50 base pair sequencing was performed on the Illumina HiSeq. The sequencing reads were aligned using HISAT2 to the UCSC dm6 genome with the options: --max-intronlen 5000 --rna-strandness F^45^. The aligned reads were then quantified using featureCounts with the Ensembl dm6 gene annotation file using the following options: -s 1 -t exon -g gene_id^46^. HISAT2 alignment statistics, PCR duplication rate (Samtools markdup^47^), and number of detected transcripts are included in Supplementary Table 7. Differential expression and principal component analysis were performed using DESeq2^48^. The genomic origin of RT&Tag reads was determined using QualiMap RNA-Seq QC^49^. IgG normalized MSL2 RT&Tag signal was visualized over the Drosophila chromosomes using karyotypeR^50^. GO term enrichment analysis for H3K27me3 and m6A enriched or depleted transcripts was performed using clusterProfiler^51^. The distribution of RT&Tag reads across the gene bodies of Drosophila genes was calculated using RSeQC^52^. For CUT&Tag, paired-end 25 base pair sequencing was performed on the Illumina HiSeq and data was analyzed as described prior (dx.doi.org/10.17504/protocols. io.bjk2kkye)^10^. MSL2 and H3K27me3 peaks were called using SEACR using the norm setting^53^. Profile plots, heatmaps and correlation matrices were generated using deepTools^54^. RRACH motifs were identified using the FIMO tool from the MEME suite^55^. Motif enrichment within the promoters of m6A enriched vs depleted transcripts was performed using the MEME tool from the MEME suit using the differential enrichment mode^56^. Genome browser screenshots were obtained from the University of California Santa Cruz (UCSC) Genome Browser. Graphs were plotted using R Studio (https://www.r-project.org) using base graphics or using ggplot2 (https://ggplot2.tidyverse.org).

## Data availability

All primary sequencing data have been deposited as single-end or paired-end fastq files in the Gene Expression Omnibus under accession code GSE195654.

## Code availability

Custom code for identifying and analyzing RT&Tag enriched transcripts is available at https://github.com/nadiyakhyzha/RTTag_Analysis.

## Acknowledgements

We thank T. Llagas and D. Xu for help with cell culture, C. Codomo for sequencing library pooling, and J. Henikoff and M. Fitzgibbon for preparing the sequencing data for analysis. This work was supported by the Howard Hughes Medical Institute (S.H.), and the Natural Sciences and Engineering Research Council (NSERC) Postdoctoral Fellowship (N.K.). The funders had no role in study design, data collection and analysis, decision to publish or preparation of the manuscript.

## Contributions

N.K., S.H., and K.A. designed the study. N.K. performed experiments and performed data analysis. N.K., S.H., and K.A. wrote the manuscript. All authors read and approved the final manuscript.

---

**Supplementary Figure 1.**
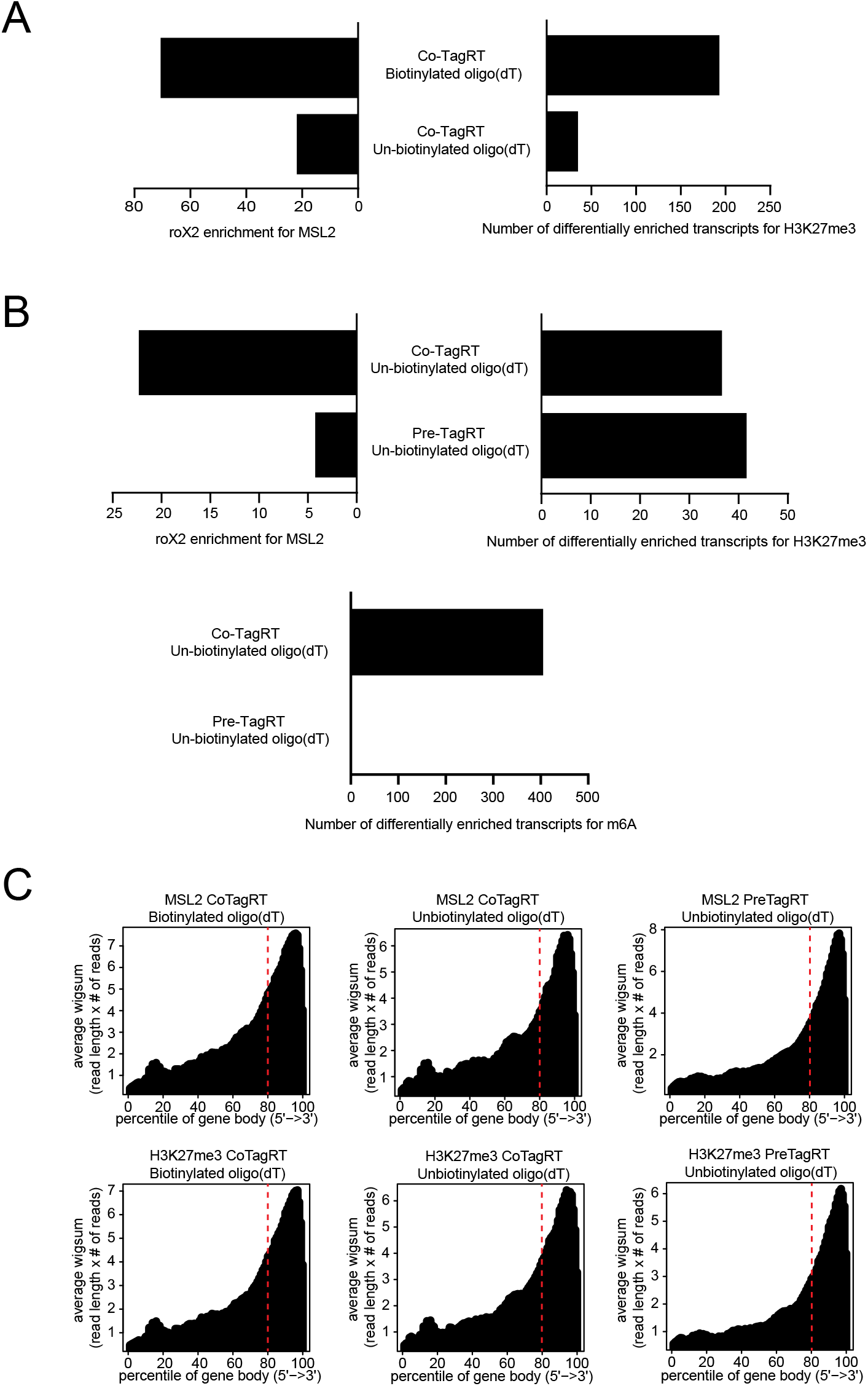
Optimization of RT&Tag. **A)** Performance comparison of RT&Tag using biotinylated or un-biotinylated oligo(dT)-adaptor B fusion oligonucleotides based on the following metrics: *roX2* enrichment for MSL2 (left) and number of differentially enriched transcripts for K27me3 (right). Both experiments were performed using reverse transcription performed at the same time as tagmentation (CoTagRT) approach. **B)** Performance comparison of RT&Tag if reverse transcription is performed prior to addition of pA-Tn5 (preTagRT) or if reverse transcription is performed at the same time as tagmentation (CoTagRT). Both experiments were performed using un-biotinylated oligo(dT)-adaptor B fusion oligonucleotides. Performance of RT&Tag was assessed based on the following metrics: *roX2* enrichment for MSL2 (top left), number of differentially enriched transcripts for K27me3 (top right) and number of differentially enriched transcripts for m6A (bottom) with pre-TagRT versus Co-TagRT. Differential enrichment was defined as >2-fold change for K27me3 or >1.5-fold change for m6A, <0.05 FDR. **C)** Density plots showing the distribution of aligned MSL2 (top) and H3K27me3 (bottom) RT&Tag reads (n=2) scaled over Drosophila gene bodies for biotinylated oligo(dT) CoTagRT (left), unbiotinylated oligo(dT) CoTagRT (center), and unbiotinylated oligo(dT) preTagRT (right) RT&Tag variations. A clear bias towards the 3’ end of genes is observed under all conditions.

**Supplementary Figure 2.**
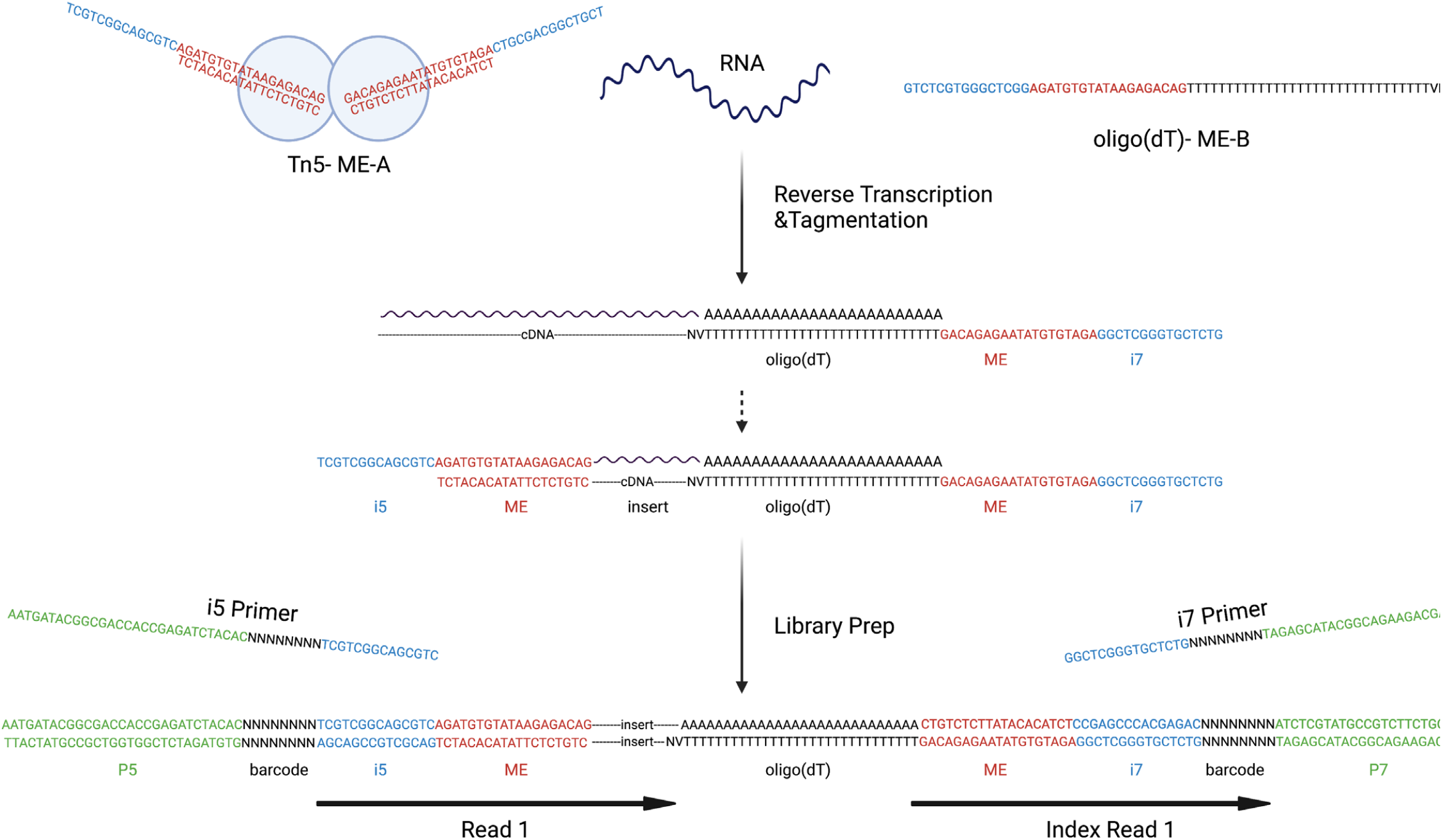
Construction of RT&Tag libraries. Schematic showing how RT&Tag libraries are generated. During reverse transcription the oligo(dT)-ME-B fusion oligonucleotide binds to the poly(A) tail of RNA. Anchored oligo(dT) is used to ensure binding at the start of the poly(A) tail. Through the process of reverse transcription, the ME-B sequence gets appended to the cDNA. The RNA/cDNA hybrid then gets tagmented with ME-A loaded Tn5. Sequencing libraries are then amplified using primers complementary to the i5 and i7 sequences. The libraries are sequenced using 50 base pair single-end sequencing with the read originating from the i5 side.

**Supplementary Figure 3.**
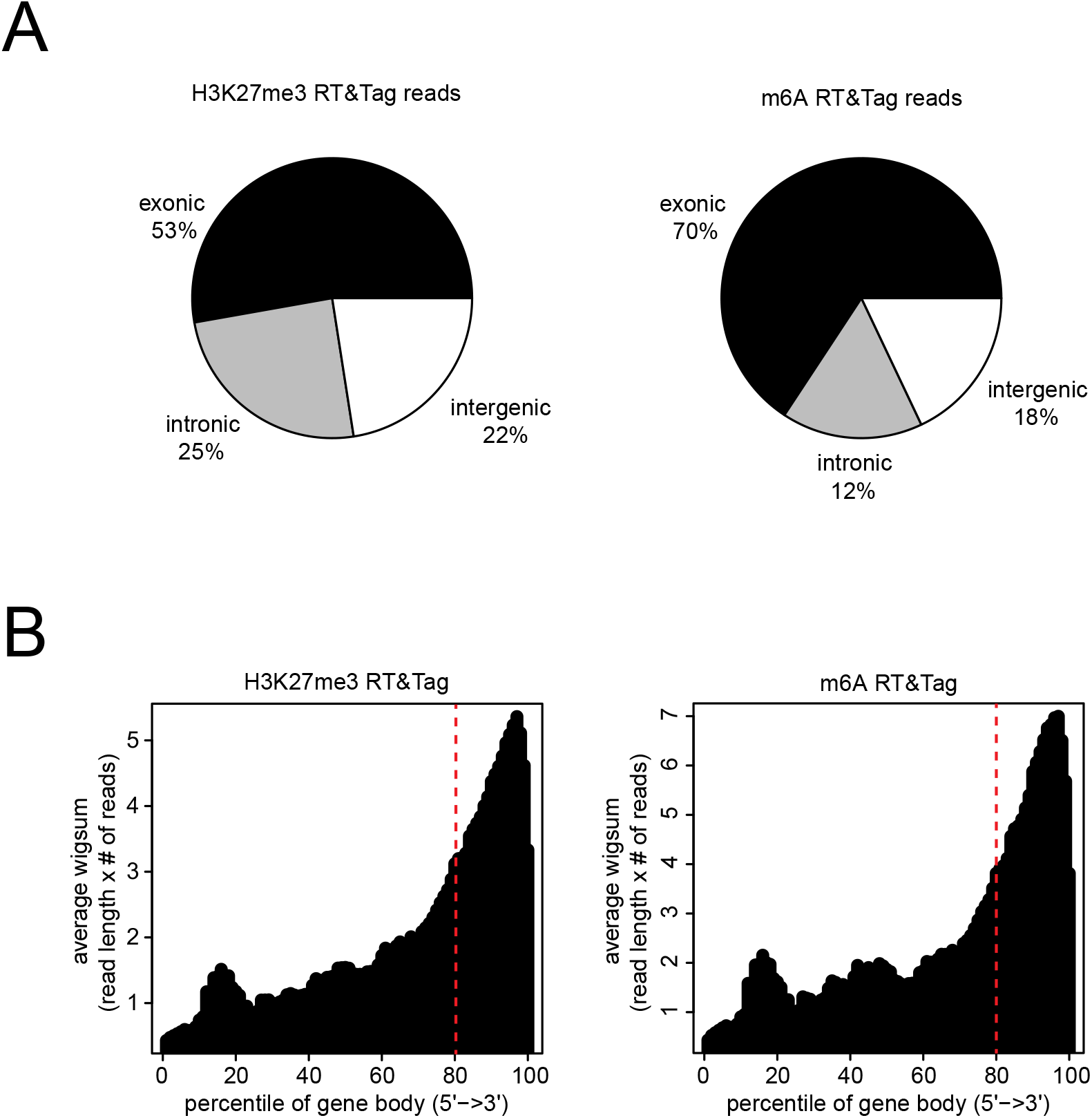
H3K27me3 and m6A RT&Tag signal. **A)** Pie chart showing the proportion of H3K27me3 (left, n=5) and m6A (right, n=3) RT&Tag reads aligning to regions classified as either exonic, intronic, or intergenic. B) Density plots showing the distribution of aligned H3K27me3 (left, n=5) and m6A (right, n=3) RT&Tag reads scaled over Drosophila gene bodies.

**Supplementary Figure 4.**
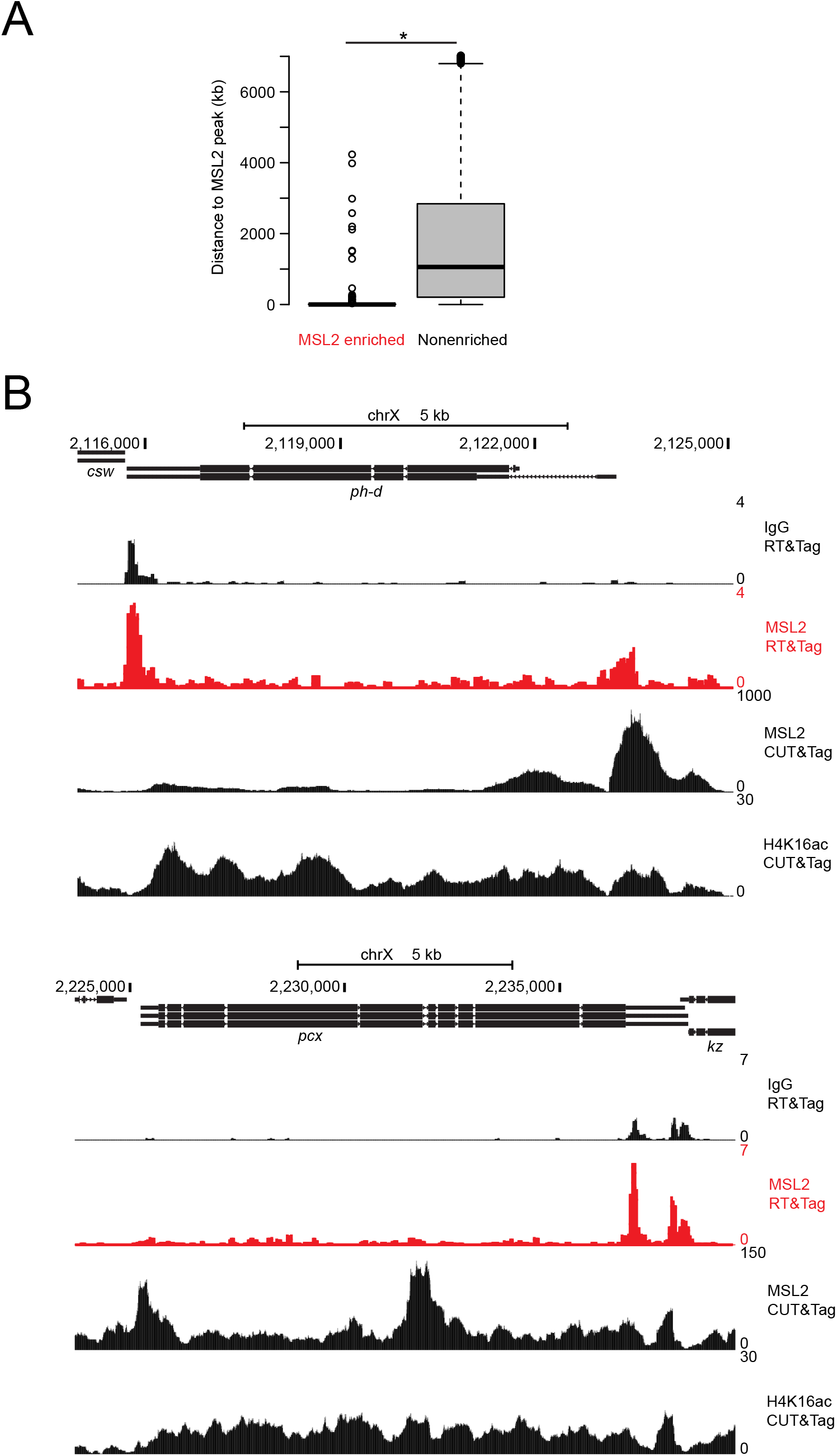
RT&Tag captures the interaction between MSL2 and transcripts within its vicinity. **A)** Boxplot showing the genomic distance from the gene body of MSL2 enriched or nonenriched transcripts to the nearest MSL2 peak. *p<0.05. **B)** Genome browser tracks showing the distribution of IgG and MSL2 RT&Tag signal as well as MSL2 and H4K16ac CUT&Tag signal over the *ph-d* and *pcx* gene bodies.

**Supplementary Figure 5.**
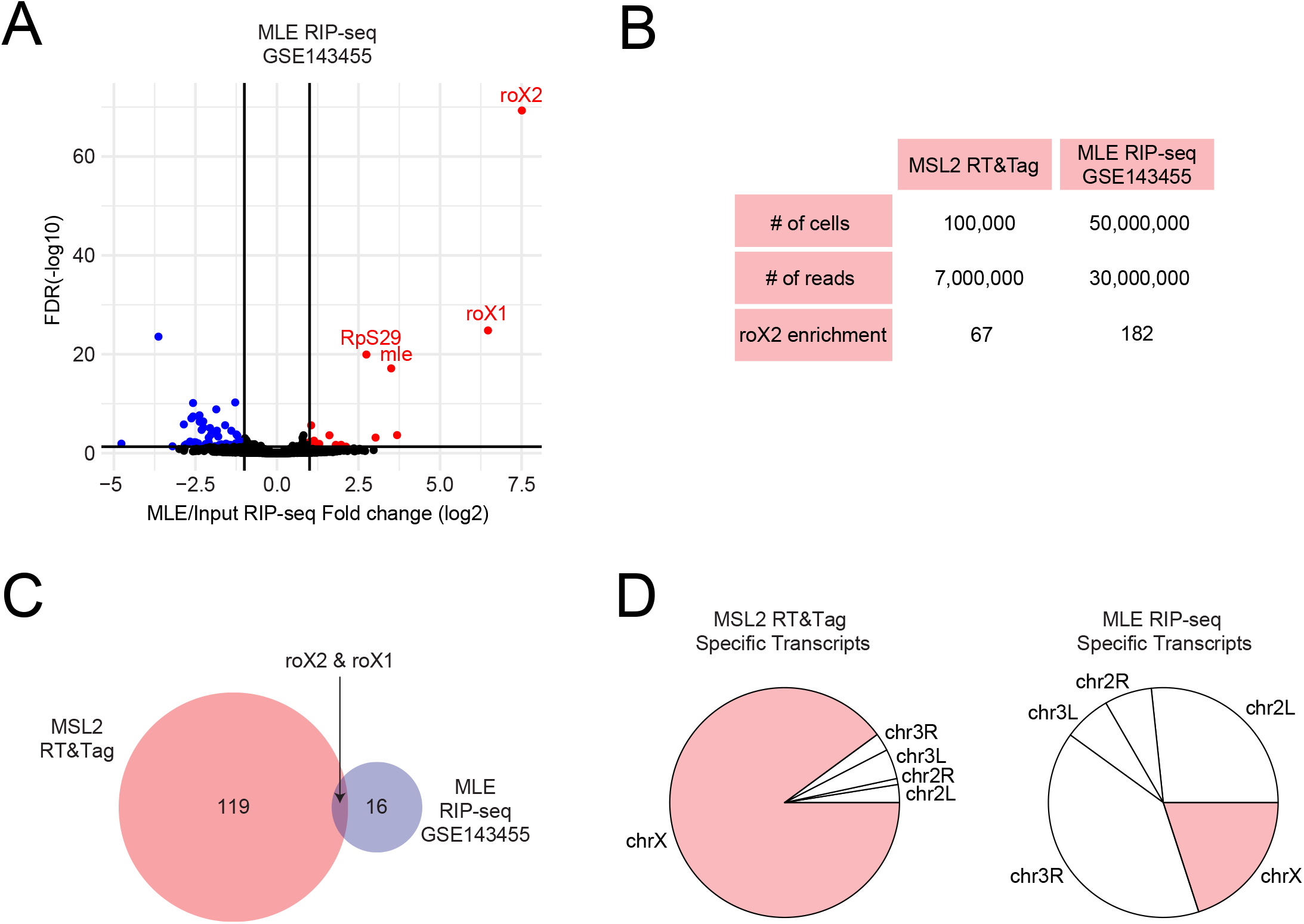
Performance comparison of MSL2 RT&Tag to RIP-seq. **A)** Volcano plot showing transcripts differentially enriched for MLE RIP-seq over input (fold change >2, FDR <0.05, n=3, GSE143455). Transcripts enriched for MLE are highlighted in red, nonenriched are in black, and depleted are in blue. **B)** Table comparing MSL2 RT&Tag and MLE RIP-seq in terms of number of cells, number of reads, and *roX2* fold change enrichment for MSL2/MLE over control. **C)** Venn diagram showing the overlap between transcripts enriched for MSL2 RT&Tag and MLE RIP-seq with *roX1* and *roX2* being enriched in both. **D)** Pie charts showing the chromosomal distribution of transcripts uniquely enriched for MSL2 RT&Tag (left) and MLE RIP-seq (right).

**Supplementary Figure 6.**
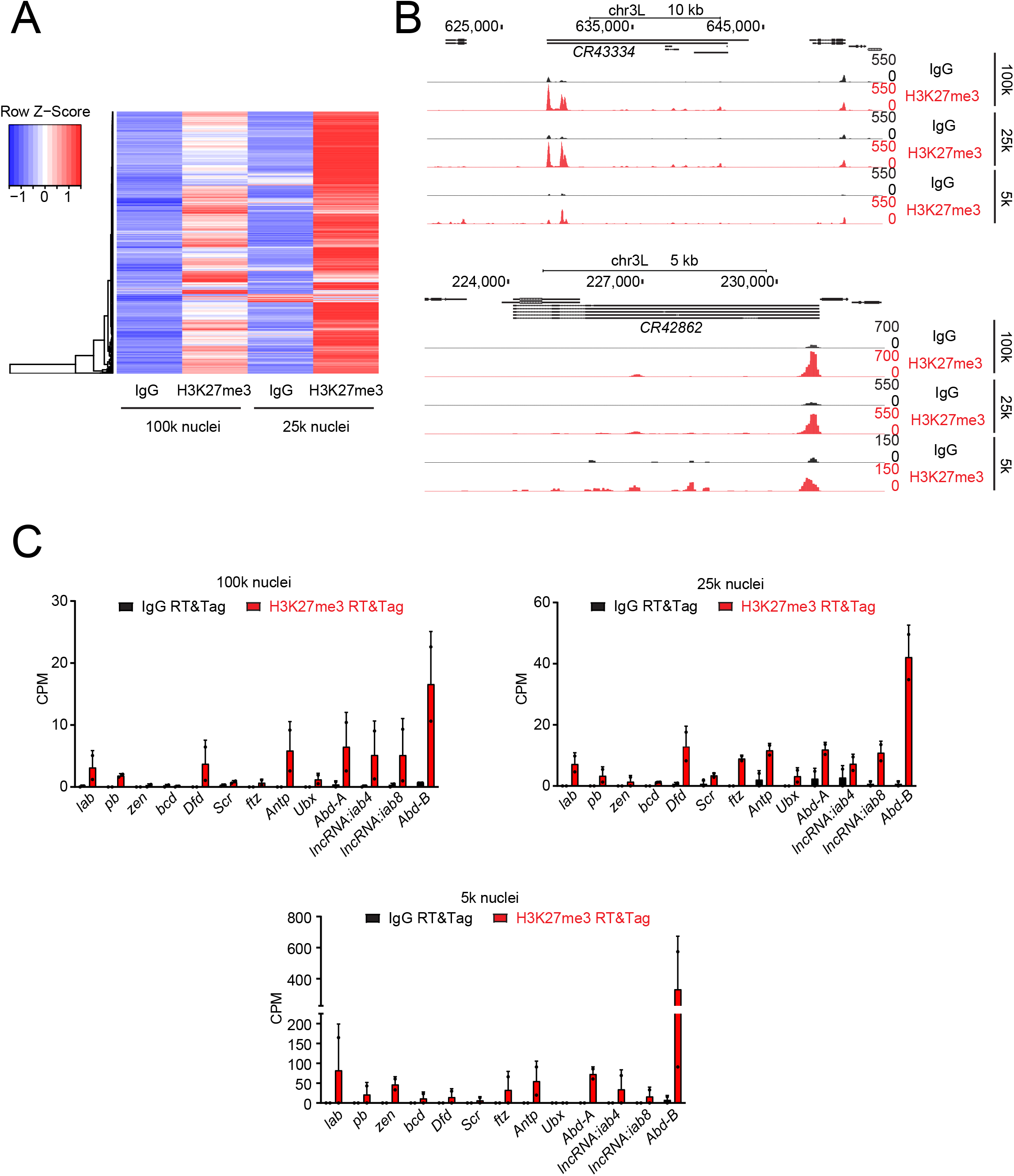
H3K27me3 RT&Tag performance with decreasing number of nuclei input. **A)** Heatmap showing the average IgG and H3K27me3 RT&Tag signal from two experiments performed using either 100,000 or 25,000 nuclei. Individual rows represent the 1342 transcripts identified as H3K27me3-enriched in Figure 3B. **B)** Genome browser tracks showing the distribution of IgG and H3K27me3 RT&Tag signal from 100,000, 25,000, or 5000 nuclei over the gene bodies of *CR43334* and *CR42862*. Combined reads from 2 replicates are shown. **C)** Boxplots showing the IgG and H3K27me3 RT&Tag signal (Counts per million, CPM) from 100,000, 25,000, or 5000 nuclei for the HOX cluster genes.

**Supplementary Figure 7.**
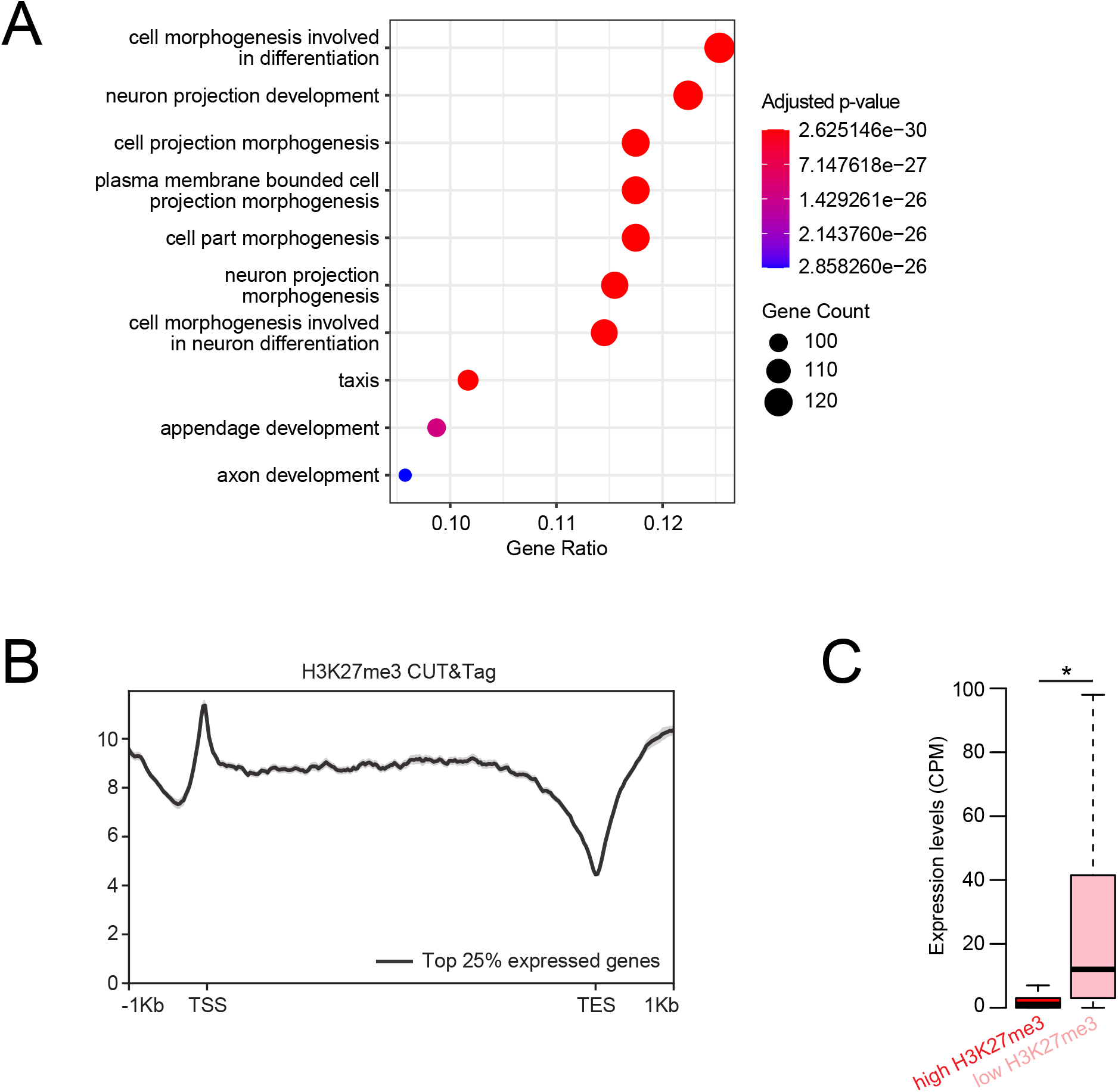
RT&Tag captures transcripts within Polycomb domains. **A)** Dot plot showing the top 10 GO biological process terms associated with H3K27me3-enriched transcripts. The dot size corresponds to the gene count and the color represents statistical significance. **B)** Profile plot showing the H3K27me3 CUT&Tag signal over the gene bodies of the top 25% expressed genes. **C)** Boxplot showing the RNA-seq expression levels (Counts per million, CPM) of H3K27me3-RT&Tag enriched transcripts that had either high (>9 read counts) or low (<9 read counts) H3K27me3 CUT&Tag signal over their gene bodies. *p<0.05, unpaired t-test.

**Supplementary Figure 8.**
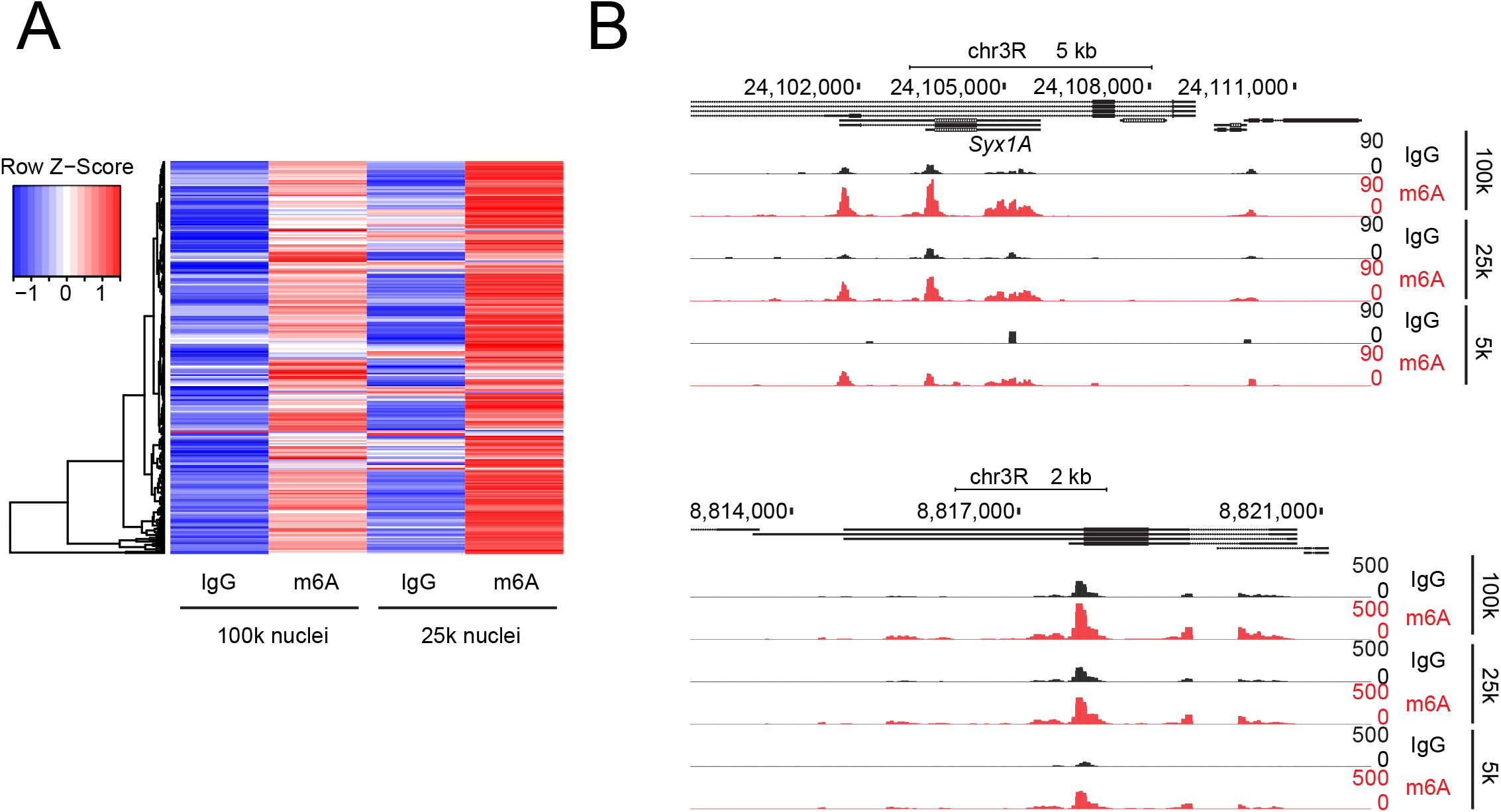
M6A RT&Tag performance with decreasing number of nuclei input. **A)** Heatmap showing the average IgG and m6A RT&Tag signal from two experiments performed using either 100,000 or 25,000 nuclei. Individual rows represent the 281 transcripts identified as m6A-enriched in Figure 4B. **B)** Genome browser tracks showing the distribution of IgG and m6A RT&Tag signal from 100,000, 25,000, or 5000 nuclei over the gene bodies of *aqz* and *Syx1A*. Combined reads from 2 replicates are shown.

**Supplementary Figure 9.**
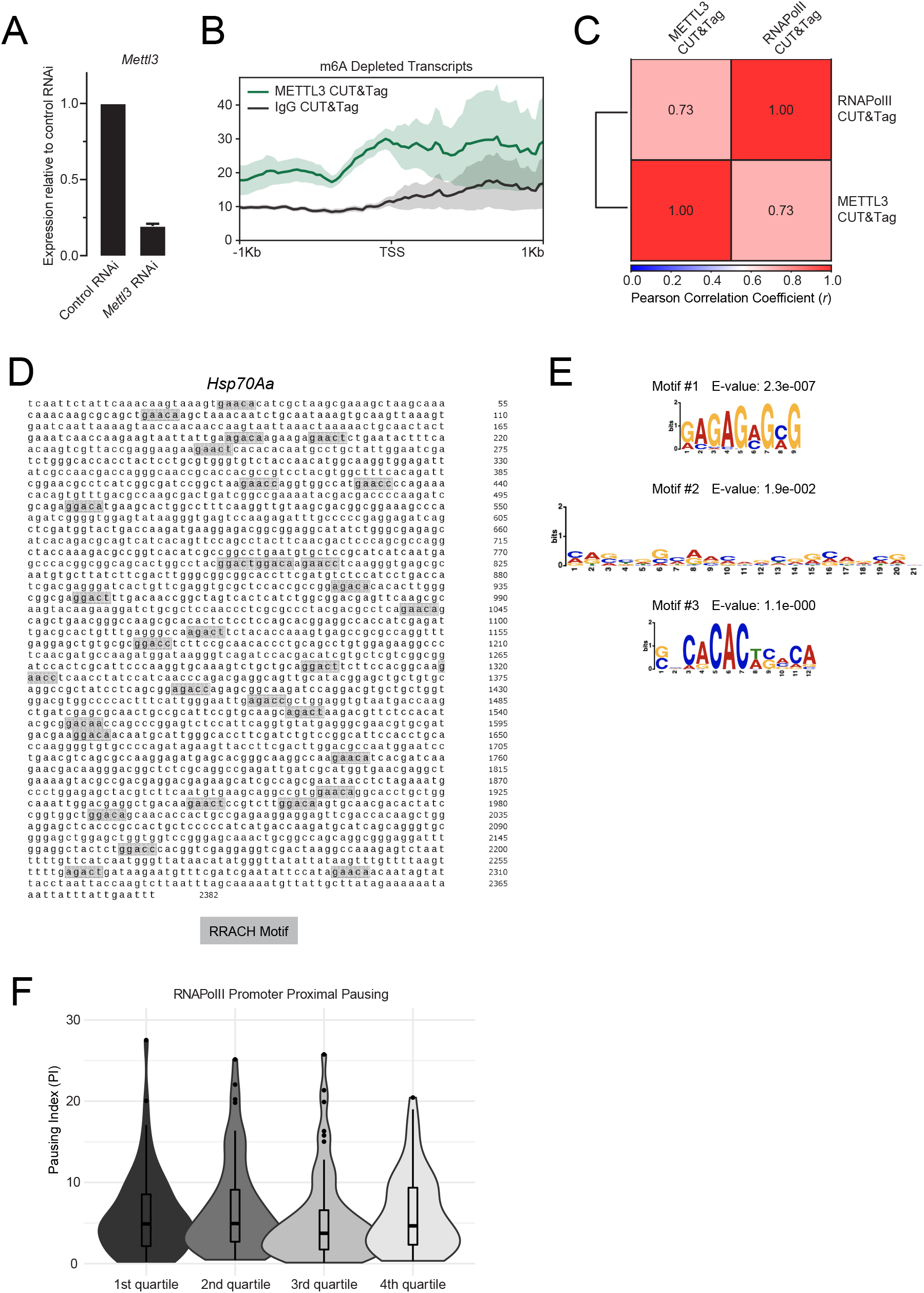
Genes of methylated transcripts are characterized by promoter proximally paused RNA Polymerase II. **A)** Bar plot showing *Mettl3* expression measured by real time PCR in control RNAi and *Mettl3* RNAi S2 cells. Data is plotted relative to control RNAi. **B)** Profile plot showing IgG and METTL3 CUT&Tag signal over the gene bodies of m6A depleted genes. **C)** Pearson correlation between RNAPolII and METTL3 CUT&Tag signal at the promoters of top 25% expressed genes. **D)** Sequence of *Hsp70Aa* with RRACH motifs highlighted in grey. **E)** MEME motif logos found to be enriched within the promoters of m6A enriched transcripts relative to those of m6A depleted transcripts using the differential enrichment mode setting. **F)** Violin plots displaying the promoter proximal pausing index (PI) m6A-enriched transcripts broken down into quartiles based on their RNA-seq expression levels. PI was calculated by dividing the promoter (+/- 250 bp around the TSS) RNAPolII CUT&Tag signal over the gene body RNAPolII CUT&Tag signal.

**Supplementary Table 1. MSL2 RT&Tag differentially enriched transcripts**.

**Supplementary Table 2. H3K27me3 RT&Tag differentially enriched transcripts**.

**Supplementary Table 3. M6A RT&Tag differentially enriched transcripts**.

**Supplementary Table 4. RT&Tag oligonucleotides**.

**Supplementary Table 5. RNAi construct primers**.

**Supplementary Table 6. Real time-PCR primers**.

**Supplementary Table 7. RT&Tag data quality statistics**.

## References

1 Licatalosi, D. D. & Darnell, R. B. RNA processing and its regulation: global insights into biological networks. Nat Rev Genet 11, 75–87, doi:10.1038/nrg2673 (2010).

2 Gagliardi, M. & Matarazzo, M. R. RIP: RNA Immunoprecipitation. Methods Mol Biol 1480, 73–86, doi:10.1007/978-1-4939-6380-5_7 (2016).

3 Zhao, J. et al. Genome-wide identification of polycomb-associated RNAs by RIP-seq. Mol Cell 40, 939–953, doi:10.1016/j.molcel.2010.12.011 (2010).

4 Moran, V. A., Niland, C. N. & Khalil, A. M. Co-Immunoprecipitation of long noncoding RNAs. Methods Mol Biol 925, 219–228, doi:10.1007/978-1-62703-011-3_15 (2012).

5 Fang, J. et al. PIRCh-seq: functional classification of non-coding RNAs associated with distinct histone modifications. Genome Biol 20, 292, doi:10.1186/s13059-019-1880-3 (2019).

6 Mondal, T., Subhash, S. & Kanduri, C. Chromatin RNA Immunoprecipitation (ChRIP). Methods Mol Biol 1689, 65–76, doi:10.1007/978-1-4939-7380-4_6 (2018).

7 Dominissini, D. et al. Topology of the human and mouse m6A RNA methylomes revealed by m6A-seq. Nature 485, 201–206, doi:10.1038/nature11112 (2012).

8 Meyer, K. D. et al. Comprehensive analysis of mRNA methylation reveals enrichment in 3’ UTRs and near stop codons. Cell 149, 1635–1646, doi:10.1016/j.cell.2012.05.003 (2012).

9 McHugh, C. A., Russell, P. & Guttman, M. Methods for comprehensive experimental identification of RNA-protein interactions. Genome Biol 15, 203, doi:10.1186/gb4152 (2014).

10 Kaya-Okur, H. S. et al. CUT&Tag for efficient epigenomic profiling of small samples and single cells. Nat Commun 10, 1930, doi:10.1038/s41467-019-09982-5 (2019).

11 Di, L. et al. RNA sequencing by direct tagmentation of RNA/ DNA hybrids. Proc. Natl. Acad. Sci. U. S. A. 117, 2886–2893, doi:10.1073/pnas.1919800117 (2020).

12 Lu, B. et al. Transposase-assisted tagmentation of RNA/DNA hybrid duplexes. Elife 9, doi:10.7554/eLife.54919 (2020).

13 Conrad, T. & Akhtar, A. Dosage compensation in Drosophila melanogaster: epigenetic fine-tuning of chromosome-wide transcription. Nat Rev Genet 13, 123–134, doi:10.1038/nrg3124 (2012).

14 Cheutin, T. & Cavalli, G. The multiscale effects of polycomb mechanisms on 3D chromatin folding. Crit Rev Biochem Mol Biol 54, 399–417, doi:10.1080/10409238.2019.1679082 (2019).

15 Blackledge, N. P. & Klose, R. J. The molecular principles of gene regulation by Polycomb repressive complexes. Nat Rev Mol Cell Biol 22, 815–833, doi:10.1038/s41580-021-00398-y (2021).

16 Lee, T. I. et al. Control of developmental regulators by Polycomb in human embryonic stem cells. Cell 125, 301–313, doi:10.1016/j.cell.2006.02.043 (2006).

17 Kassis, J. A., Kennison, J. A. & Tamkun, J. W. Polycomb and Trithorax Group Genes in Drosophila. Genetics 206, 1699–1725, doi:10.1534/genetics.115.185116 (2017).

18 Bell, J. C. et al. Chromatin-associated RNA sequencing (ChAR-seq) maps genome-wide RNA-to-DNA contacts. Elife 7, doi:10.7554/eLife.27024 (2018).

19 Li, X. et al. GRID-seq reveals the global RNA-chromatin interactome. Nat. Biotechnol. 35, 940–950, doi:10.1038/nbt.3968 (2017).

20 He, P. C. & He, C. m(6) A RNA methylation: from mechanisms to therapeutic potential. EMBO J 40, e105977, doi:10.15252/embj.2020105977 (2021).

21 McIntyre, A. B. R. et al. Limits in the detection of m(6)A changes using MeRIP/m(6)A-seq. Sci Rep 10, 6590, doi:10.1038/s41598-020-63355-3 (2020).

22 Kan, L. et al. A neural m(6)A/Ythdf pathway is required for learning and memory in Drosophila. Nat Commun 12, 1458, doi:10.1038/s41467-021-21537-1 (2021).

23 Lence, T., Soller, M. & Roignant, J. Y. A fly view on the roles and mechanisms of the m(6)A mRNA modification and its players. RNA Biol 14, 1232–1240, doi:10.1080/15476286.2017.1307484 (2017).

24 Haussmann, I. U. et al. m(6)A potentiates Sxl alternative pre-mRNA splicing for robust Drosophila sex determination. Nature 540, 301–304, doi:10.1038/nature20577 (2016).

25 Akhtar, J. et al. m(6)A RNA methylation regulates promoter-proximal pausing of RNA polymerase II. Mol Cell 81, 3356–3367 e3356, doi:10.1016/j.molcel.2021.06.023 (2021).

26 Guertin, M. J., Petesch, S. J., Zobeck, K. L., Min, I. M. & Lis, J. T. Drosophila heat shock system as a general model to investigate transcriptional regulation. Cold Spring Harb Symp Quant Biol 75, 1–9, doi:10.1101/sqb.2010.75.039 (2010).

27 Chetverina, D., Erokhin, M. & Schedl, P. GAGA factor: a multifunctional pioneering chromatin protein. Cell Mol Life Sci 78, 4125–4141, doi:10.1007/s00018-021-03776-z (2021).

28 Pallares, L. F., Picard, S. & Ayroles, J. F. TM3’seq: A Tagmentation-Mediated 3’ Sequencing Approach for Improving Scalability of RNAseq Experiments. G3 (Bethesda) 10, 143–150, doi:10.1534/g3.119.400821 (2020).

29 Fazal, F. M. et al. Atlas of Subcellular RNA Localization Revealed by APEX-Seq. Cell 178, 473–490 e426, doi:10.1016/j.cell.2019.05.027 (2019).

30 Padron, A., Iwasaki, S. & Ingolia, N. T. Proximity RNA Labeling by APEX-Seq Reveals the Organization of Translation Initiation Complexes and Repressive RNA Granules. Mol Cell 75, 875–887 e875, doi:10.1016/j.molcel.2019.07.030 (2019).

31 McMahon, A. C. et al. TRIBE: Hijacking an RNA-Editing Enzyme to Identify Cell-Specific Targets of RNA-Binding Proteins. Cell 165, 742–753, doi:10.1016/j.cell.2016.03.007 (2016).

32 Janssens, D. H. et al. Automated CUT&Tag profiling of chromatin heterogeneity in mixed-lineage leukemia. Nat Genet 53, 1586–1596, doi:10.1038/s41588-021-00941-9 (2021).

33 Muniz, L., Nicolas, E. & Trouche, D. RNA polymerase II speed: a key player in controlling and adapting transcriptome composition. EMBO J 40, e105740, doi:10.15252/embj.2020105740 (2021).

34 Slobodin, B. et al. Transcription Impacts the Efficiency of mRNA Translation via Co-transcriptional N6-adenosine Methylation. Cell 169, 326–337 e312, doi:10.1016/j.cell.2017.03.031 (2017).

35 Lence, T. et al. m(6)A modulates neuronal functions and sex determination in Drosophila. Nature 540, 242–247, doi:10.1038/nature20568 (2016).

36 Xu, W. et al. Dynamic control of chromatin-associated m(6) A methylation regulates nascent RNA synthesis. Mol Cell, doi:10.1016/j.molcel.2022.02.006 (2022).

37 Louloupi, A., Ntini, E., Conrad, T. & Orom, U. A. V. Transient N-6-Methyladenosine Transcriptome Sequencing Reveals a Regulatory Role of m6A in Splicing Efficiency. Cell Rep 23, 3429–3437, doi:10.1016/j.celrep.2018.05.077 (2018).

38 Van Nostrand, E. L. et al. A large-scale binding and functional map of human RNA-binding proteins. Nature 583, 711–719, doi:10.1038/s41586-020-2077-3 (2020).

39 Gerstberger, S., Hafner, M. & Tuschl, T. A census of human RNA-binding proteins. Nat Rev Genet 15, 829–845, doi:10.1038/nrg3813 (2014).

40 Gebauer, F., Schwarzl, T., Valcarcel, J. & Hentze, M. W. RNA-binding proteins in human genetic disease. Nat Rev Genet 22, 185–198, doi:10.1038/s41576-020-00302-y (2021).

41 Geula, S. et al. Stem cells. m6A mRNA methylation facilitates resolution of naive pluripotency toward differentiation. Science 347, 1002–1006, doi:10.1126/science.1261417 (2015).

42 Li, H. B. et al. m(6)A mRNA methylation controls T cell homeostasis by targeting the IL-7/STAT5/SOCS pathways. Nature 548, 338–342, doi:10.1038/nature23450 (2017).

43 Lee, H. et al. Stage-specific requirement for Mettl3-dependent m(6)A mRNA methylation during haematopoietic stem cell differentiation. Nat Cell Biol 21, 700–709, doi:10.1038/s41556-019-0318-1 (2019).

44 Batista, P. J. et al. m(6)A RNA modification controls cell fate transition in mammalian embryonic stem cells. Cell Stem Cell 15, 707–719, doi:10.1016/j.stem.2014.09.019 (2014).

45 Kim, D., Paggi, J. M., Park, C., Bennett, C. & Salzberg, S. L. Graph-based genome alignment and genotyping with HISAT2 and HISAT-genotype. Nat Biotechnol 37, 907–915, doi:10.1038/s41587-019-0201-4 (2019).

46 Liao, Y., Smyth, G. K. & Shi, W. featureCounts: an efficient general purpose program for assigning sequence reads to genomic features. Bioinformatics 30, 923–930, doi:10.1093/bioinformatics/btt656 (2014).

47 Li, H. et al. The Sequence Alignment/Map format and SAMtools. Bioinformatics 25, 2078–2079, doi:10.1093/bioinformatics/btp352 (2009).

48 Love, M. I., Huber, W. & Anders, S. Moderated estimation of fold change and dispersion for RNA-seq data with DESeq2. Genome Biol 15, 550, doi:10.1186/s13059-014-0550-8 (2014).

49 Okonechnikov, K., Conesa, A. & Garcia-Alcalde, F. Qualimap 2: advanced multi-sample quality control for high-throughput sequencing data. Bioinformatics 32, 292–294, doi:10.1093/bioinformatics/btv566 (2016).

50 Gel, B. & Serra, E. karyoploteR: an R/Bioconductor package to plot customizable genomes displaying arbitrary data. Bioinformatics 33, 3088–3090, doi:10.1093/bioinformatics/btx346 (2017).

51 Yu, G., Wang, L. G., Han, Y. & He, Q. Y. clusterProfiler: an R package for comparing biological themes among gene clusters. OMICS 16, 284–287, doi:10.1089/omi.2011.0118 (2012).

52 Wang, L., Wang, S. & Li, W. RSeQC: quality control of RNAseq experiments. Bioinformatics 28, 2184–2185, doi:10.1093/bioinformatics/bts356 (2012).

53 Meers, M. P., Tenenbaum, D. & Henikoff, S. Peak calling by Sparse Enrichment Analysis for CUT&RUN chromatin profiling. Epigenetics Chromatin 12, 42, doi:10.1186/s13072-019-0287-4 (2019).

54 Ramirez, F. et al. deepTools2: a next generation web server for deep-sequencing data analysis. Nucleic Acids Res 44, W160–165, doi:10.1093/nar/gkw257 (2016).

55 Grant, C. E., Bailey, T. L. & Noble, W. S. FIMO: scanning for occurrences of a given motif. Bioinformatics 27, 1017–1018, doi:10.1093/bioinformatics/btr064 (2011).

56 Bailey, T. L. & Elkan, C. Fitting a mixture model by expectation maximization to discover motifs in biopolymers. Proc Int Conf Intell Syst Mol Biol 2, 28–36 (1994).

